# Genomic islands of divergence reveal selection on the alternate homeolog in upper Fraser River sockeye run-timing groups

**DOI:** 10.1101/2025.06.04.657877

**Authors:** Ben J. G. Sutherland, Cory Williamson, Brian Toth, Gordon Sterritt

## Abstract

Effective wildlife conservation and management depends on a thorough understanding of both neutral and adaptive intraspecific genomic variation. Although population structure is often well understood for many ecologically or economically important species, the extensive population genomic resources required to characterize putatively adaptive variation, for example in punctuated genomic regions, is typically only recently available in many species, if at all. Here we explore recently published whole-genome resequencing data in the upper Fraser River and identify a large island of divergence separating populations with different run timings in the Nechako Watershed. The island of divergence at approximately 56.3-58.0 Mbp of Chr18 has extended homozygosity and reduced Tajima’s D in specific populations of the upper Fraser River. These include the Nadina and Stellako River sockeye, as well as the Stuart-Summer run timing group, but not the geographically proximal Early Stuart sockeye. In the populations without the Stuart-Summer haploblock, genetic variation looks more similar to elsewhere throughout the chromosome. When investigating the gene content of this region, importantly it was determined to contain the *leucine rich-repeat containing 9-like* (*lrrc9-like*) gene and surrounding homeologous region to the *lrrc9* region of Chr12, known to contain a conserved and selected island of divergence that has been associated with life history traits and run timing throughout the species range. By also inspecting the samples in the present work at the Chr12 haploblock, three genotypic states were observed throughout the region, although the Stuart River run timing groups contained mainly the reference haplotype. Using similar methods, no other regions of major differentiation were observed in the available samples for the Chilcotin or Quesnel River systems. These results provide additional evidence for the importance of both homeologous regions holding *lrrc9* and *lrrc9-like* in different geographic regions and underscore the uniqueness of upper Fraser River sockeye in this potentially adaptive island of divergence.

## Introduction

Genetic differentiation provides insights into population distinctiveness (Weir and Cockerham 1984) and can have significant implications for conservation and species management. It can be characterized with 10s of neutral microsatellite markers (e.g., Beacham et al. 2004), 100s of single nucleotide polymorphism (SNP) markers by amplicon panels (Meek and Larson 2019), 10,000s of SNPs by reduced representation sequencing (Baird et al. 2008; Peterson et al. 2012) or 100,000s to millions of SNPs through whole-genome resequencing (Christensen et al. 2024; Bemmels et al. 2025). Although these approaches can recover similar trends, they can also differ due to the inherent difference in sampling breadth across the genome and the presence of punctuated genomic regions (e.g., Barry et al. 2024) that can involve adaptive variation (Tigano and Russello 2022). These regions can also involve structural variation such as inversions (Akopyan et al. 2025) that can prevent local recombination and keep haplotypes largely intact over time, providing an adaptive benefit if holding co-adapted allele complexes (Wellenreuther and Bernatchez 2018).

Over the past decade significant discoveries have identified major effect loci associated with run timing and other reproductive strategies in salmonids. Alleles differentiating migration timing groups have been identified in the GREB1L/ROCK1 region in both steelhead trout *Oncorhynchus mykiss* and Chinook salmon *O. tshawytscha* (Hess et al. 2016; Prince et al. 2017; Thompson et al. 2019; Thompson et al. 2020; Narum et al. 2024). Although the orthologous region is involved in both species, the putatively adaptive variation evolved independently (Prince et al. 2017). This can occur if there is a limited number of pathways to the mechanism, the trait is relatively simple (e.g., monogenic control), or if the genetic background incurring selection has shared variation (Wood et al. 2005). These discoveries have prompted the significant question as to whether these types of loci should be specifically considered in conservation efforts (Pearse 2016; Waples et al. 2022). This is a critical question, for example given the ecological and economic value of the portfolio effect (the buffering that comes with intraspecific variation; Schindler et al. 2010) as well as that specific run timing forms can become locally extirpated or rarer from differential anthropogenic pressures (e.g., Anderson et al. 2025).

Other putative major effect loci have been identified in the salmonids. In Atlantic salmon *Salmo salar*, the *vgll3*/*six6* region was found on Chr25 and is associated with control of age-at-maturity (Barson et al. 2015; Sinclair-Waters et al. 2022). Chr25 is the ancestral salmonid chromosome 16.2 using the naming convention outlined by Sutherland et al. (2016). In sockeye salmon *O. nerka*, islands of divergence have been identified in sockeye salmon throughout the past decade, with regions of major differentiation being observed in association with different life history types (Larson et al. 2017; Veale and Russello 2017; Larson et al. 2019; Tigano and Russello 2022; Euclide et al. 2024). Although these often do not have consistent range-wide phenotypic associations, one exception is a region on Chr12 (salmonid ancestral 15.1) near the gene *leucine rich repeat-containing 9* (*lrrc9*). This region has been associated with beach or stream spawning range-wide (Veale and Russello 2017; Tigano and Russello 2022) and run timing (Larson et al. 2017; Larson et al. 2019; Barry et al. 2025). Recently, it was discovered that the orthologous region is also involved in run timing in chum salmon *O. keta*, and in pink salmon *O. gorbuscha* and coho salmon *O. kisutch* (Barry et al. 2025). Moreover, in chum and coho salmon, the homeologous region (i.e., on salmonid ancestral 15.2) also showed an island of divergence associated with run timing (Barry et al. 2025). In all the different species cases, the putative adaptation appears to have independently evolved (Barry et al. 2025), as was described for the GREB1L/ROCK1 region (Prince et al. 2017). Mechanisms remain largely unknown in these studies, but major effect loci appear to use a “relatively simple” genetic system to produce “remarkable diversity” (Anderson et al. 2025) that is essential for the portfolio effect (Schindler et al. 2010) and to allow for adaptation to local ecological niches (Lisi et al. 2013).

The Fraser River watershed in British Columbia (BC), Canada is the largest sockeye salmon complex in Canada, and has the greatest sockeye abundance in the world for any single river system (COSEWIC 2017). Over the past century, there have been significant declines in all five species of salmon in the Fraser River (Northcote and Atagi 1997). Genetic evidence suggests that among the Pacific salmon, sockeye have had particularly drastic declines (Christensen et al. 2024). Multiple salmon species show similar trends in hierarchical population genetic structure in the Fraser River, with the greatest differences observed among broad regional groupings (i.e., lower, middle, and upper) (Christensen et al. 2024). The overall population structure of sockeye salmon in the Fraser has been documented using multiple different marker types, including microsatellites and SNPs (Beacham et al. 2004; Rondeau 2022). Subgroupings and structure among spawning locations is generally based on geographic proximity as well as ecotype and run timing (Beacham et al. 2004; Beacham and Withler 2017). The genetic diversity of Fraser River salmon collectively represents the standing genetic variation for the species in the region and holds adaptive traits and genetic diversity that is irreplaceable on timescales that are relevant to humans (Waples et al. 2001; COSEWIC 2017) critical for fisheries, culture, and resilience to climate change.

A major sub-basin of the Fraser River watershed in the upper-most regions of the system is the upper Stuart River Watershed including Takla Lake and Trembler Lake (Bradford and Braun 2021). This watershed contains different run-timing groups with relatively close geographic spawning locations: Early Stuart sockeye, the sole representative of the ‘Early Stuart’ run timing group; and Stuart-Summer sockeye, a genetically distinct run timing group (Beacham et al. 2004; Rondeau 2022). Both the Early Stuart and Stuart-Summer sockeye are classified as endangered populations (COSEWIC 2017). Early Stuart sockeye have the earliest upstream migration of all Fraser River sockeye (Idler and Clemens 1959), with returning adults entering the Fraser River in early July then quickly migrating to spawning streams for peak spawning in mid-August (Bradford and Braun 2021). Approximately half of the Early Stuart run timing group passes Hell’s Gate by July 14^th^, whereas the early summer, summer, and late run timing groups correspond to Aug. 6^th^, Aug. 17^th^, and Sept. 21^st^, respectively (COSEWIC 2017). The Stuart River Watershed sockeye must travel great distances and overcome significant river force to reach the spawning grounds (Wright et al. 2025). This is thought to have shaped Early Stuart sockeye genetics and physiology through long-exerted evolutionary forces based on energetic requirements and local environments of the route (Hinch and Rand 1998; Crossin et al. 2004; Rand et al. 2006; Eliason et al. 2011). For example, Early Stuart sockeye have smaller and more fusiform body shapes than other deeper bodied morphologies that occur in sockeye populations with easier routes, as well as higher somatic energy density, fewer carried eggs, and more efficient travel (Gilhousen 1980; reviewed by Crossin et al. 2004), and have the capability for higher aerobic scope and cardiac capacity (Crossin et al. 2004; Eliason et al. 2011). The Early Stuart metapopulation (Bradford and Braun 2021), comprised of over 50 spawning sites (COSEWIC 2017), has not been observed to have genetic differentiation between streams as evaluated by microsatellite or amplicon panels (Beacham et al. 2004; Rondeau 2022). However, a more thorough investigation of this genome-wide is merited to support decisions around hatchery broodstock crosses (e.g., Bradford et al. 2025) in the recently developed Early Stuart sockeye conservation hatcheries (UFFCA 2026).

The present study aims to characterize at the whole-genome level the variation that exists within sockeye salmon in the upper Fraser River to support population conservation including hatchery management. It is also of significant conservation interest to determine whether any genomic regions (novel or previously identified elsewhere) are associated with different run-timing phenotypes in the region. Run timing is expected to provide resilience to climate change through its portfolio effect (Schindler et al. 2010; Anderson et al. 2015) and the different run timing groups are expected to be differentially impacted by the effects of climate change (Rand et al. 2006; Reed et al. 2011). An understanding of the presence and uniqueness of any putative functional genetic variation related to this trait within the region is therefore of high importance in developing conservation approaches range-wide. By using recently published SNP data from whole-genome resequencing of sockeye salmon (Christensen et al. 2024), we directly address these two main objectives in the upper Fraser River.

## Methods

### Samples and genotypes

Multilocus genotypes were obtained as a VCF file from a recent study on salmonids of the Fraser River (see Data Availability; Christensen et al. 2024). Samples were originally sourced from genetic baseline collections of returning adults to spawning grounds from DFO (Christensen et al. 2024). The VCF file was subset with bcftools (Danecek et al. 2021) to retain sockeye collections from the mid- or upper-Fraser River (n = 210 samples; Table 1; Figure 1). Although some variation in year sampled occurs (Table 1), temporal genetic differentiation was found to be negligible relative to spatial variation for these populations, in particular at the timescales investigated here (Beacham et al. 2004). Age of adults (return age) was not available for the samples within metadata, other than that the sockeye were returning adults at or near spawning grounds (i.e., baseline collection). Following the sample subset, allele frequencies were recalculated, and only those SNPs with a minor allele frequency (MAF) > 0.01 were retained. The SNP positions in the VCF file were from an unpublished genome assembly used by Christensen et al. (2024) (K. Christensen, *pers. comm.*), and so these were converted to the latest genome available in NCBI (i.e., GCF_034236695.1; Oner_Uvic_2.0; Christensen et al. 2020) using SNPlift (Normandeau et al. 2023). An additional dataset was generated using bcftools for use in exploratory population genetic analyses by filtering for MAF > 0.05 and pruning SNPs for linkage disequilibrium (LD; maximum r^2^ = 0.5 in 50 kb windows).

**Figure 1.**
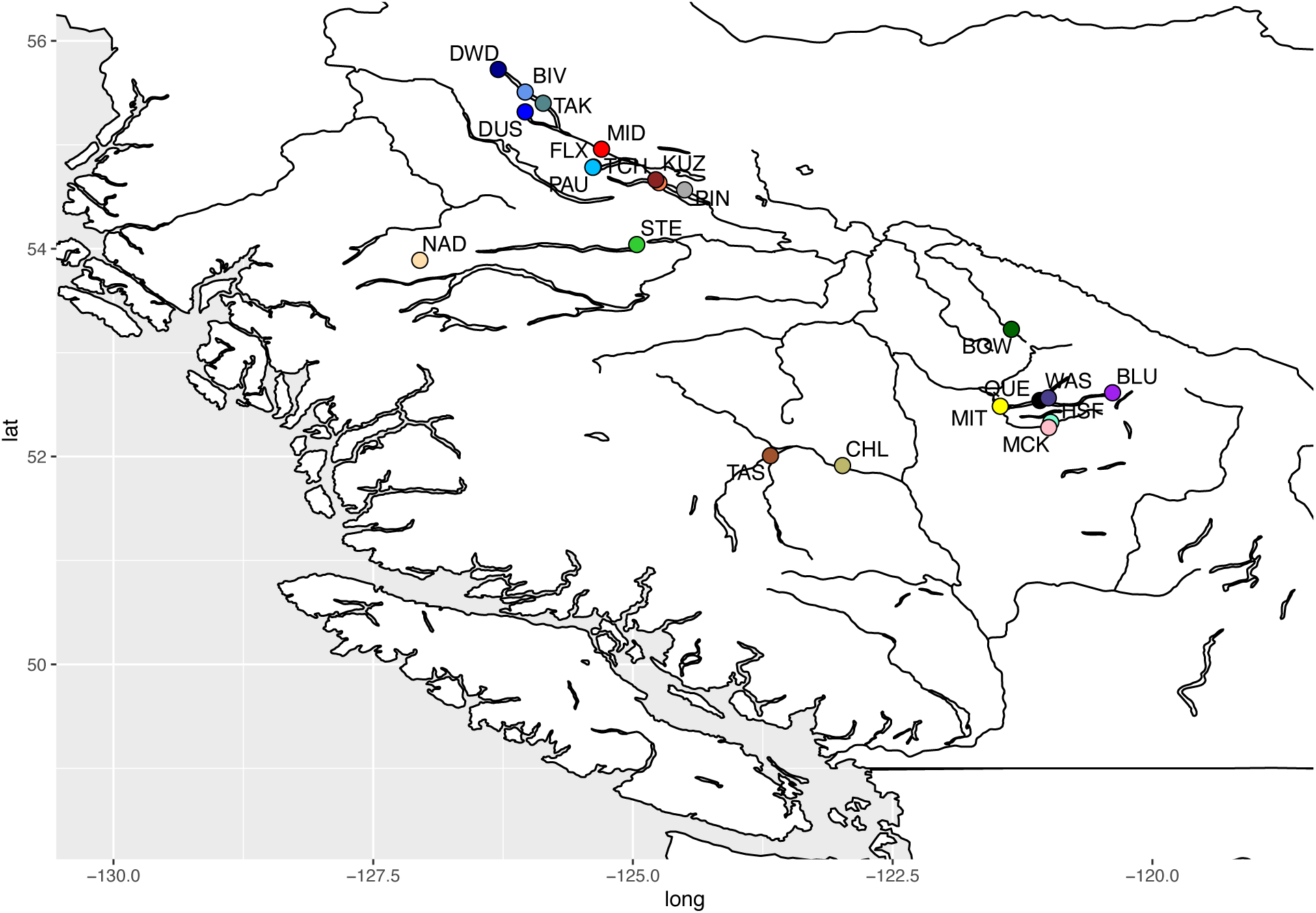
Locations of sockeye salmon populations used for whole-genome resequencing in the upper Fraser River of British Columbia. Samples were from the Early Stuart metapopulation (DWD=Driftwood R.; BIV=Bivouac Cr.; TAK=Takla Lk.; DUS=Dust Cr.; FLX=Felix Cr.; PAU=Paula Cr.), Stuart-Summer populations (MID=Middle R.; TCH=Tachie R.; KUZ=Kuzkwa R.; PIN=Pinchie Cr.), Nadina R. (NAD), Stellako R. (STE), Bowron R. (BOW), Quesnel Lake Watershed (QUE=Quesnel Lk.; MIT=Mitchell Bay; WAS=Wasko Cr.; BLU=Blue Lead Cr.; HSF=Horsefly R.; and MCK=McKinley Cr.), and the Chilcotin Watershed (CHL=Chilcotin R.; TAS=Taseko R.). More details on populations are provided in Table 1.

**Table 1.** Samples available for the analysis from Christensen et al. (2024) genotyped by whole-genome resequencing. Collection names and dates were provided in sample metadata from the original study, full names were determined from BC Geographical Names (Government of BC), and run-timing designations were obtained from Rondeau (2022).

| Region | Run-Timing | Collection | Full Name | Sample size | Year(s) sampled |
| --- | --- | --- | --- | --- | --- |
| Stuart | Early-Stuart | Bivouac | Bivouac Cr. | 10 | 2005 |
|  |  | Driftwood | Driftwood R. | 7 | 2005 |
|  |  | Dust | Dust Cr. | 10 | 2005 |
|  |  | Felix | Sidney (Felix) Cr. | 5 | 2005 |
|  |  | Paula | Paula Cr. | 10 | 2005 |
|  |  | Takla | Takla Lk. | 4 | *unkn.* |
|  | Summer | Kuzkwa | Kuzkwa R. | 9 | 2001 |
|  |  | Middle | Middle R. | 7 | 2001 |
|  |  | Pinchi | Pinchi Cr. | 4 | 2005 |
|  |  | Tachie | Tachie R. | 11 | 2012, *unkn.* |
| Nadina/Francois | Early summer | Nadina | Nadina R. | 9 | 2000 |
| Francois/Fraser | Summer | Stellako | Stellako R. | 10 | 2011 |
| Bowron | Early summer | Bowron | Bowron R. | 10 | 2001 |
| Quesnel | Summer | BlueLead | Blue Lead Cr. | 5 | 2001 |
|  |  | McKinley | McKinley Cr. | 9 | 2001 |
|  |  | Mitchell | Mitchell Bay | 8 | 2001 |
|  |  | Wasko | Wasko Cr. | 5 | 2001 |
|  |  | Horsefly* | Horsefly R. and Lk. | 24 | 2001, *unkn* |
|  |  | Quesnel | Quesnel R. and Lk. | 8 | 2013 |
| Chilcotin | Early Summer | Taseko | Taseko R. | 10 | 2011 |
|  | Summer | Chilko* | Chilko R. and Lk. | 35 | 2001, 2008, *unkn* |
|  |  | TOTAL |  | 210 |  |

\*includes various sub-naming

### Population genetics

The MAF > 0.05 and LD-filtered VCF file was read into R (R Core Team 2026) using vcfR (Knaus and Grünwald 2017), converted to a genind object using *vcf2genind* of vcfR, then converted to genlight object using *gi2gl* of dartR (Gruber et al. 2018). The genlight object was used in a principal components analysis (PCA) including all samples using *glPca* of adegenet (Jombart and Ahmed 2011). The genind object was also used to construct a dendrogram with 10,000 bootstraps with poppr (Kamvar et al. 2014) using the Edwards distance metric (Cavalli-Sforza and Edwards 1967). Subsequent region-specific PCA analyses were also conducted as above, with datasets filtered for samples from collections from (1) all waters that flow into the Nechako River (i.e., Early Stuart, Stuart-Summer, Nadina River, and Stellako River); (2) all waters that flow into the Quesnel River (i.e., Quesnel and Horsefly); and (3) all waters that flow into the Chilcotin River (i.e., Chilko Lake, Chilko River, Taseko River; Figure 1). All sample metadata was obtained from Christensen et al. (2024), and the map locations used here are only approximations that are based on the collection site names. A map of population locations was generated using ggplot2 (Wickham 2016) with base map data obtained from the maps package (Becker et al. 2025), additional layers provided by the Natural Earth project (naturalearthdata.com), and shapefiles prepared by maptools (Bivand and Lewin-Koh 2013). Average *F*_ST_ (Weir and Cockerham 1984) between collections was determined by calculating site-specific *F*_ST_ in specific contrasts using VCFtools (Danecek et al. 2011), then calculating 95% confidence intervals from these site-specific values in R.

To further characterize run timing groups within watersheds, the datasets described above were prepared and imported into R, but only filtered for MAF > 0.01 (and no LD filter). Per watershed, monomorphic loci were removed, and a discriminant analysis of principal components (DAPC) was run using adegenet to identify any loci that contributed substantially to separating collections. Per allele (reference allele) DAPC loading values were plotted in a Manhattan plot across sockeye salmon chromosomes using fastman (Paria et al. 2022). The same approach was taken for only the Early Stuart collections to investigate for any punctuated regions of differentiation within the metapopulation.

### Islands of divergence characterization

Based on the results from the above DAPC and the previously identified region on sockeye salmon Chr12 (Barry et al. 2025), VCF files for each of Chr18 and Chr12 were generated using bcftools. These were used for per-locus *F*_ST_ calculations using pegas (Paradis 2010) in R. Subsequently, the R package *rehh* (Gautier et al. 2017; Klassmann and Gautier 2022) was used to calculate per-population extended haplotype homozygosity using the *scan_hh* function, then the rolling average of the output integrated site-specific EHH (iES) score was determined using *rollapply* of zoo package (v.1.8.15; Zeileis and Grothendieck 2005). Results were plotted using line graphs in R. Given the EHH results, site-specific Tajima’s D (Tajima 1989) was calculated using VCFtools across Chr18 for the Early Stuart and Stuart-Summer collections at the reporting unit scale (i.e., all collections within the conservation unit).

To understand the geographic context of the observed differentiated region on Chr18, the approximate outlier section of Chr18 with *F*_ST_ > 0.5 and an elevated iES score was identified and SNPs within this region for all individuals in the study were selected using bcftools and read into R using vcfR. Genotypes were converted to allele dosage and plotted using the *heatmap* function of base R. A heatmap was generated for the Nechako region (Table 1, regions Stuart, Nadina/Francois and Francois/Fraser) with dendrogram clustering of samples, and a separate heatmap was generated for the entire dataset using collections sorted by reporting unit from north to south (Table 1).

To inspect the genic content of the differentiated region on Chr18, the identified region plotted in the heatmap above was inspected for annotated genes in NCBI. The corresponding gene content on Chr12 was obtained given strong syntenic relationships of ohnologs, and the corresponding window to the region of interest on Chr18 was determined for Chr12. Both regions and full chromosomes were then subjected to a snpEff analysis to estimate effects of the characterized SNPs (Cingolani et al. 2012) using a custom database built from the Oner_Uvic_2.0 assembly and gene annotations to determine the putative impacts of genetic variants observed. A heatmap of the corresponding region on Chr12 was generated and a scan for EHH was calculated and plotted as described for Chr18 above. Gene acronyms were obtained manually from online sources and plotted alongside *F*_ST_ between Early Stuart and Stuart-Summer populations, predicted snpEff impacts, and the approximate region of the highly linked SNPs within the haplotype blocks (as determined visually within the heatmap) on both chromosomes using karyoploteR (Gel and Serra 2017).

The genotypes previously identified to be associated with stream/early or beach/late by Veale and Russello (2017) and Barry et al., (2024) respectively, were linked to the present genotype data using several steps. First, the primers for the TaqMan assay One_LRRC9_68810 were obtained from Veale and Russello (2017) and used as a BLAST query against the Oner_Uvic_2.0 assembly, finding the forward primer aligned at 63,418,398-63,418,423 bp of LG12 and reverse primer at 63,418,483-63,418,451 bp, with the inside of the amplicon therefore contained between 63,418,424-63,418,460 bp. This short region was visually inspected against the probe sequences provided by Veale and Russello (2017), following a reverse complement of each, where the target variant is at position 63,418,449 bp of NC_088407.1 (LG12). A ‘GG’ genotype was designated by Veale and Russello (2017) as beach spawning phenotype, and ‘GT’ or ‘TT’ as stream spawning. The nucleotides were determined from the reference genome and VCF file as reference (REF) = T and alternate (ALT) = G. Given that the stream type is also termed the early allele (Barry et al. 2024), the stream/early genotype was considered in the present work as REF/REF (T/T) or REF/ALT (T/G), and the beach/late genotype as ALT/ALT (G/G) at this locus.

## Results

### Population genetic structure in upper Fraser River sockeye salmon

Upper Fraser River sockeye salmon samples (n = 210) genotyped by Christensen et al. (2024) that are included in the present analysis are presented in Table 1 and Figure 1 and include the following regions: Stuart (Early Stuart, Stuart-Summer), Nadina/Francois (Nadina River), Francois/Fraser (Stellako River), Bowron, Quesnel (and Horsefly), and Chilcotin. The Stuart, Nadina, and Stellako locations collectively are also termed the Nechako region here. For each collection included, year of sampling was obtained from Christensen et al. (2024), and run timing designations were sourced from Rondeau (2022).

Overall population genetic trends were initially explored using LD filtered SNPs (MAF > 0.05; n = 627,421 SNPs). A PCA clustering all samples by genotypes shows four main clusters in PC1 and PC2 (Figure 2A; PVE = 3.6% and 1.8%, respectively), specifically (1) all populations with natal habitat upstream to the Nechako River (i.e., both Stuart River Watershed run timing groups as well as Nadina and Stellako); (2) the Bowron River; (3) Quesnel/Horsefly Watershed; and (4) the Chilcotin Watershed (Figure 2A). Populations from Quesnel, Horsefly, and Chilcotin regions have a similar position on PC1 but are separated across PC2. A genetic dendrogram using these samples shows similar groupings, with the largest difference in the data being between the populations upstream to the Nechako from the other collections (Figure S1). The second grouping in the dendrogram contains Bowron River as the most external population in the grouping, followed by Chilcotin River Watershed populations, and a grouping containing Quesnel and Horsefly populations, which are each grouped in separate branches (Figure S1). Some indication of unexpected clustering due to low sample size is suggested by the dendrogram, with Felix (n = 5) and Takla (n = 4) collections grouping closely and separately from other collections, as well as poor clustering of Middle River (n = 7) and Pinchi (n = 4). Driftwood (n = 7) also clusters on the outside of the other Early Stuart collections.

**Figure 2.**
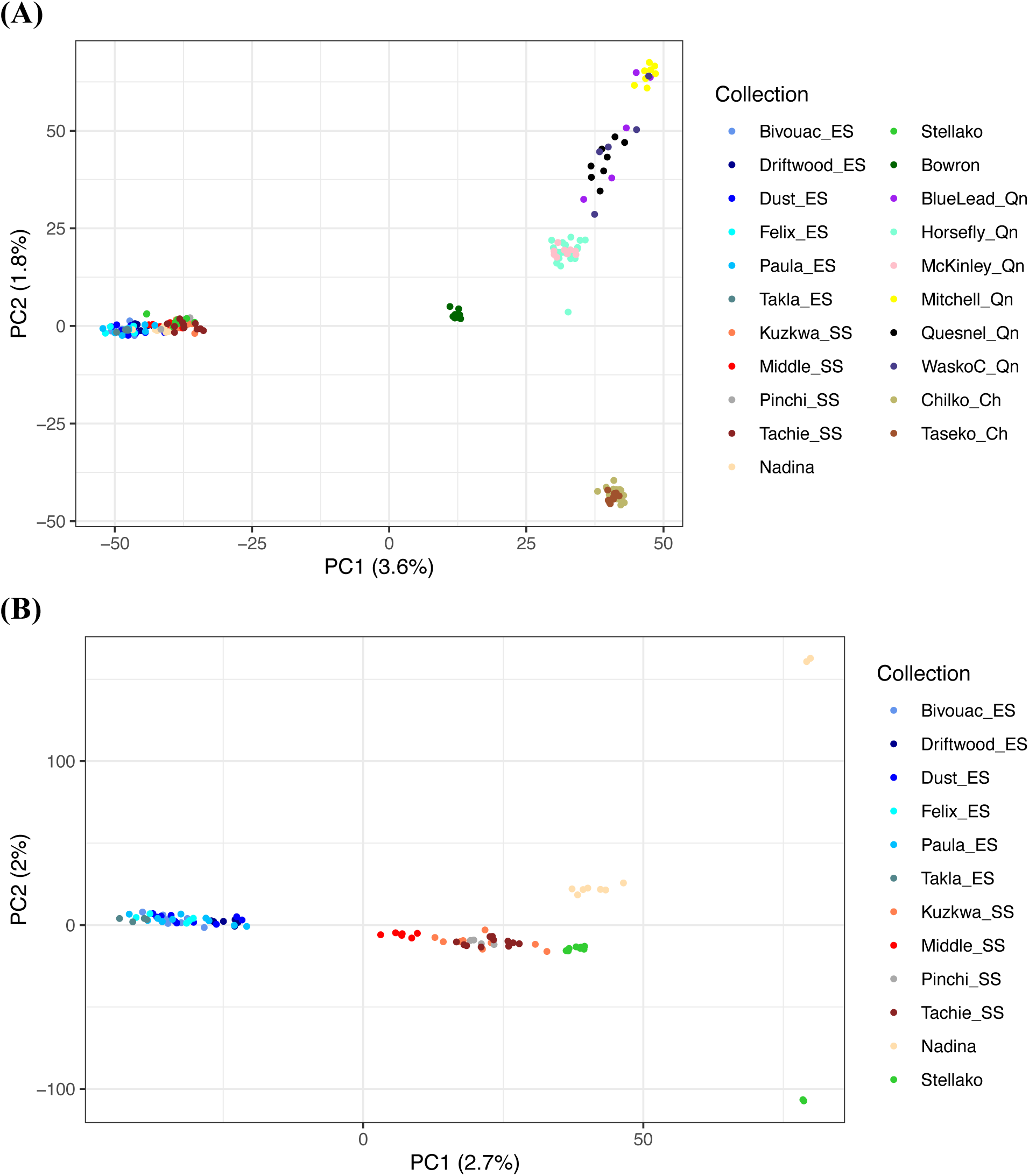
Principal components analysis (PCA) clustering samples by genotypes (MAF ≥ 0.05; LD-filtered) including samples from (A) throughout upper Fraser R.; and (B) the Nechako Watershed. The percent variation explained per axis is shown. Acronyms: ES = Early Stuart; SS = Stuart-Summer; Qn = Quesnel region; Ch = Chilcotin region.

Within-region variation was inspected for populations in the Nechako region (n = 96 samples, 12 collection sites, and 620,378 SNPs) using PCA. All Early Stuart collections grouped separately from Stuart-Summer, Nadina, and Stellako on PC1 (PVE = 2.7%; Figure 2B). Stellako River samples group closest to the Stuart-Summer collections, with Nadina River samples separated on PC2. Several outlier samples for both Nadina and Stellako River collections are observed. Applying a DAPC to these Nechako region populations (MAF > 0.01; n = 3,714,493 SNPs) produces three separate groups: (1) Early Stuart; (2) Stuart-Summer and Stellako; and (3) Nadina (Figure 3A). Plotting DAPC loading values per locus shows that loci contributing the most to this separation were clustered in a region on Chr18, showing a peak that is significantly elevated relative to the rest of the genome (Figure 3B; see below). Whereas loading values across most of Chr18 are at baseline and less than 0.00002, those in the peak are above 0.00008 (Figure 3B).

**Figure 3.**
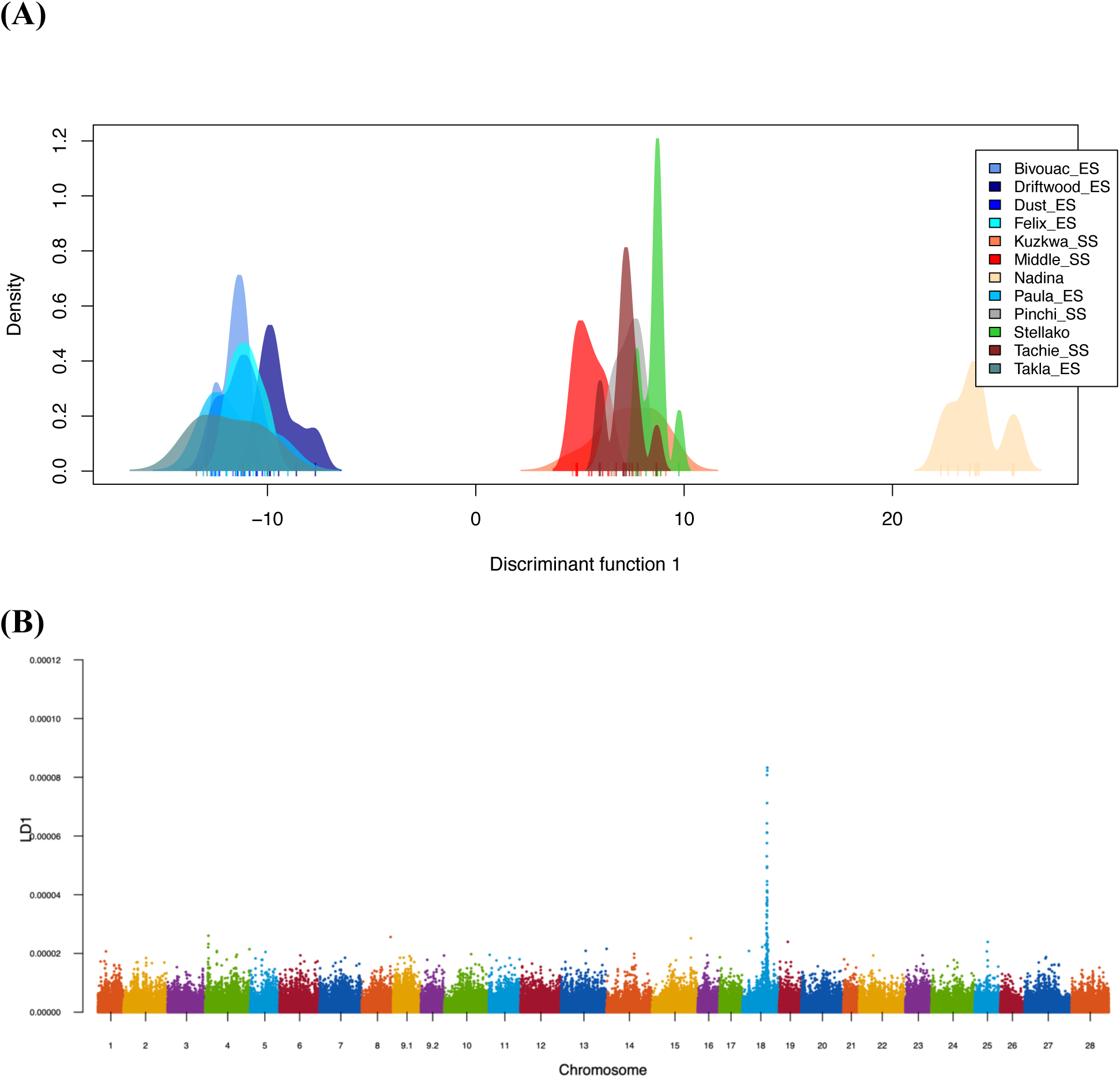
**(A)** Discriminant analysis of principal components (DAPC) of populations from the Nechako Watershed including Early Stuart (ES), Stuart-Summer (SS), Nadina R., and Stellako R. **(B)** Per-SNP loading values contributing to the DAPC separation shown in (A). Acronyms: ES = Early Stuart; SS = Stuart-Summer.

Populations upstream to the Chilcotin River (n = 45 samples, 610,932 SNPs) evaluated by PCA found that all Chilko Lake and Chilko River samples (run timing: Summer) grouped together and separately from the Taseko River collection (run timing: Early Summer; Figure S2A; PC1 PVE = 5.3%). These two groups were strongly separated by DAPC (Figure S2B; n = 3,491,171 SNPs), but the loci separating the groups came from throughout the genome without any clear outlier regions (Figure S2C). Populations upstream to the Quesnel River (n = 59 samples, 612,184 SNPs) evaluated by PCA found overlap in groupings within some Quesnel collections, and separate clustering of the Horsefly collections (i.e., Horsefly and McKinley; Figure S3A; PC1 PVE = 5.4%). A DAPC recovered similar trends (Figure S3B; n = 3,646,984 SNPs), with loci contributing to the separation of Quesnel and Horsefly found throughout the genome, with no clear indication of outlier genomic regions (Figure S3C).

### Island of divergence on Chr18 shows signal of positive selection in specific populations

The major differentiating region on Chr18 observed between the Early Stuart and the Stuart-Summer, Nadina and Stellako populations (Figure 3B; *see above*) had elevated *F*_ST_ from approximately 56.0 – 58.3 Mbp, with the highest *F*_ST_ from 57.0 – 57.5 Mbp (Figure 4A; top differentiated loci genotypes by collection in Additional File S1). Using SNPs from the region of interest and surrounding area (55.5 – 58.7 Mbp), a heatmap showed separate clustering of the above differentiated populations (Early Stuart vs. all other Nechako region populations; Figure 4B). Stuart-Summer, Nadina, and Stellako samples all showed a region of conserved homozygosity from around 57.0 – 57.9 Mbp. By contrast, the Early Stuart samples had heterozygosity observed throughout the region (conserved homozygosity not present), and by viewing this genomic region in all 210 upper Fraser River samples it was apparent that only the Stuart-Summer, Nadina, and Stellako regions hold this conserved homozygosity region. The genomic region therefore appears to be largely a homozygous conserved haplotype (haploblock) unique to Stuart-Summer, Nadina, and Stellako (Additional File S2; Table 2). A few other samples showed high homozygosity in the Quesnel Watershed and Chilcotin Watershed, but with some differences from the Stuart-Summer haploblock.

**Figure 4.**
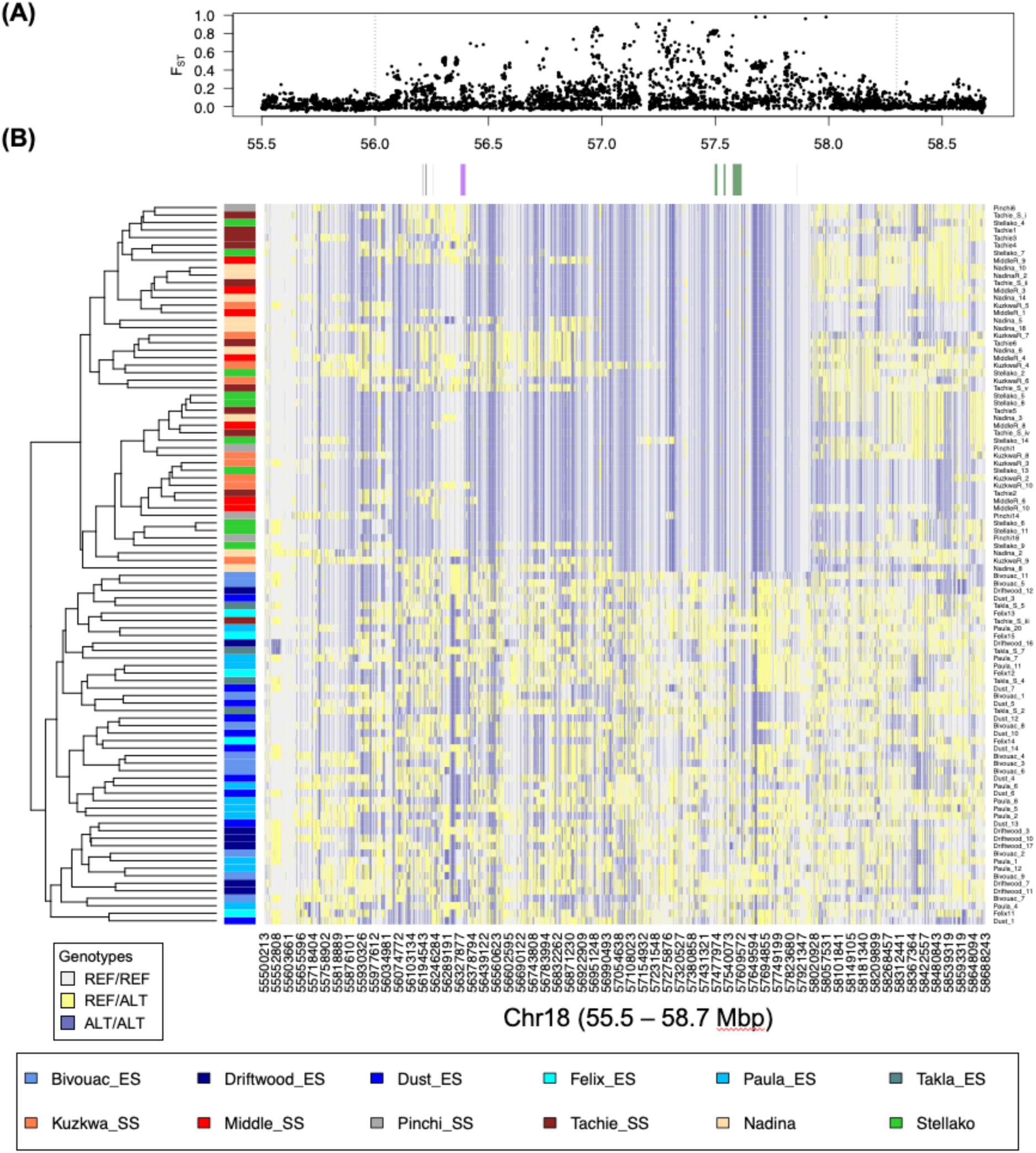
The differentiating region between 56.0-58.3 Mbp of Chr18 is shown with the immediate surrounding region for the Nechako Watershed populations. Average per-locus *F*_ST_ (A) shows significant differentiation, and the heatmap (B) clustered separately the Early Stuart metapopulation from the others in the watershed. A general lack of heterozygous genotypes within the region of interest (57.0-58.0 Mbp) are observed in the Stuart-Summer populations, as well as Stellako R., and Nadina R. sockeye. Gene positions are given above the heatmap (left to right): grey = *six6* genes; purple = *lrrc9-like*; green = *60 kDa lysophospholipase-like*; darkblue = *OTX2*. Acronyms: ES = Early Stuart; SS = Stuart-Summer.

**Table 2.**
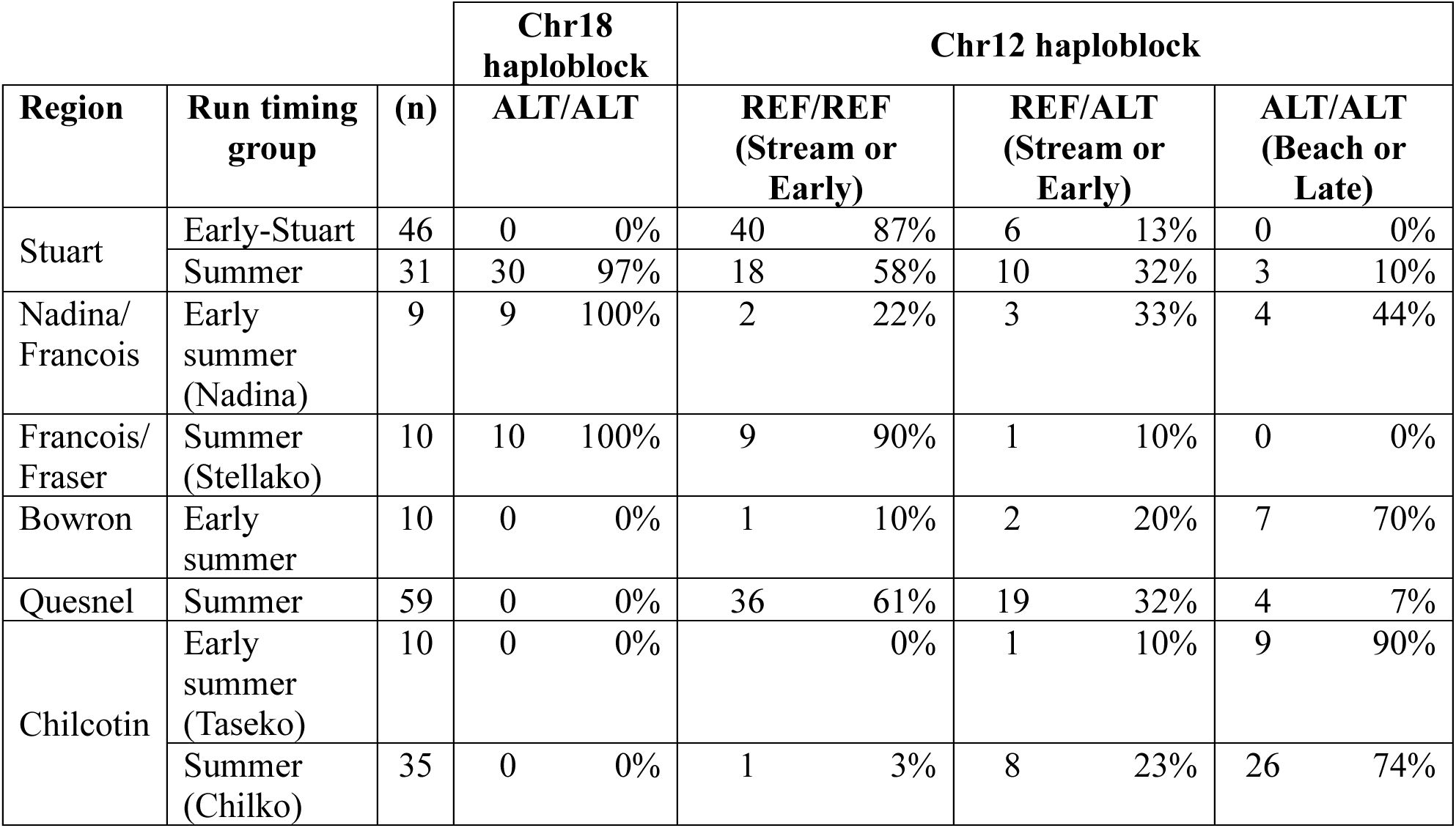
Haploblock genotype frequencies and percentages for Chr18 and Chr12 genomic regions of interest. Regional and run timing groups are shown with sample size (n). For Chr18, only the homozygous alternate haploblock is tallied, as the other genotypes do not show strong linkage. For Chr12, given the strong linkage for all haploblocks, all three genotypic haploblock states are shown.

An analysis of extended haplotype homozygosity for Chr18 indicated a strong signal for positive selection throughout the region of interest in Stuart-Summer, Nadina, and Stellako populations (Figure 5A), and not in the other populations. In agreement with this observation, Tajima’s D was significantly decreased at the island of divergence in Stuart-Summer (Figure 5C) but not Early Stuart (Figure 5D). This finding is concordant with the lack of homozygosity viewed in the heatmap for Early Stuart (Figure 4B), suggesting that only the Stuart-Summer, Nadina, and Stellako populations have evidence for positive selection at this region on Chr18.

**Figure 5.**
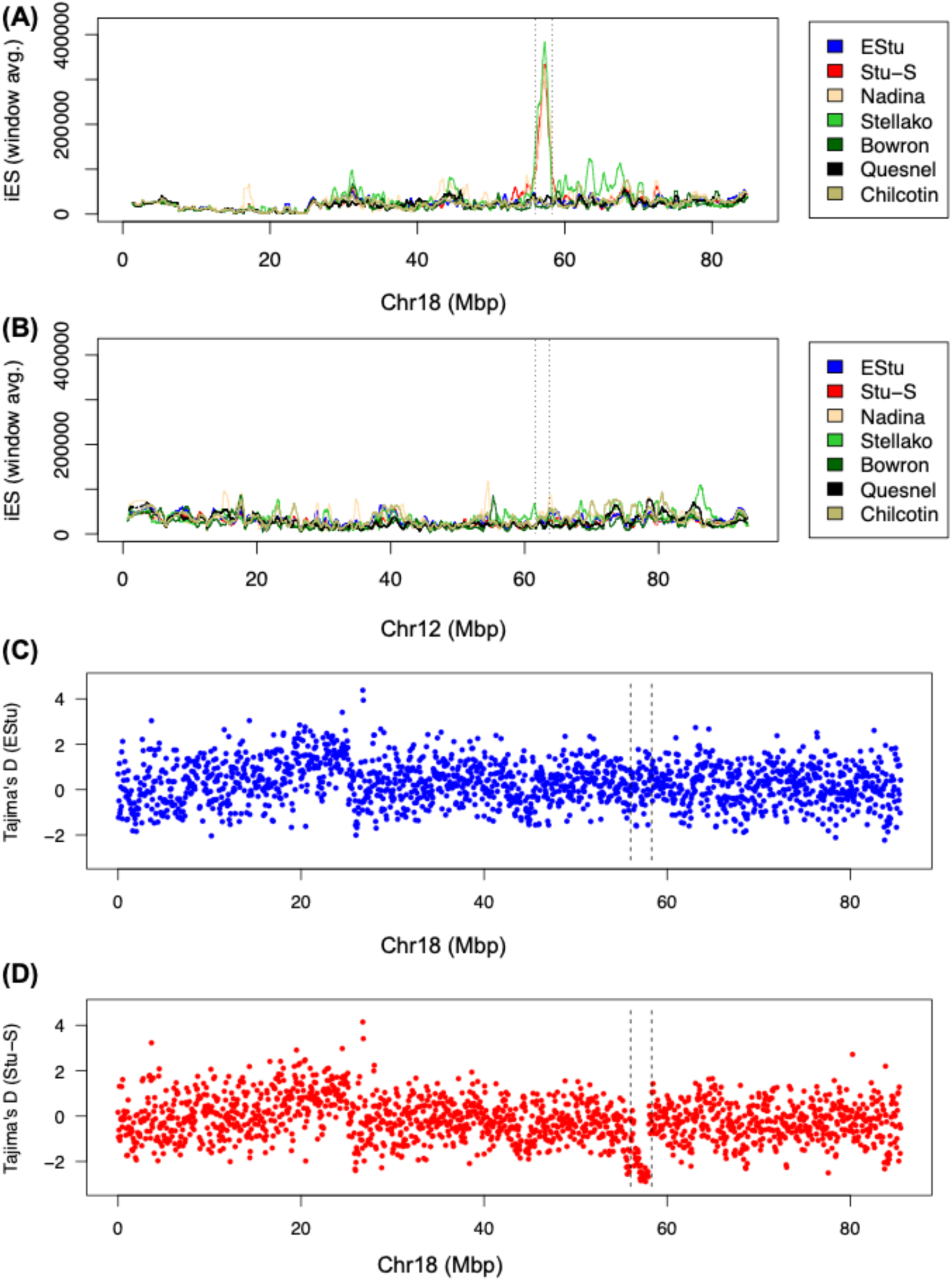
Chr18 shows evidence of positive selection as shown by (A) the integrated Extended Haplotype Homozygosity (iES), but only in Stuart-Summer, Nadina, and Stellako populations, and not the Early Stuart metapopulation or other Fraser River collections. (B) The Chr12 region does not show an elevated iES score in any collection. (C-D) Selection on Chr18 is also suggested by negative Tajima’s D values at the island of divergence in Stuart-Summer (D) but not Early Stuart samples (C). Hatched lines indicate the region of interest at 56.0-58.3 Mbp.

Gene content in the Chr18 island of divergence region (56.0 – 58.3 Mbp) included 56 protein coding genes, 9 pseudogenes, and 10 long non-coding RNA. Upon reviewing the genes in the region, the ohnolog of a gene of interest on Chr12, *leucine-rich repeat-containing protein-9* (*lrrc9*; NCBI Gene ID: 115138988), *lrrc9-like* (56.33-56.36 Mbp; NCBI Gene ID: 115146781) was observed. Interestingly, *lrrc9-like* is a predicted pseudogene and the RNA-seq aggregate read coverage (NCBI) does not match its expected exon pattern, unlike that of *lrrc9* (Figure S5). Notably, other previously identified genes related to run-timing in this region included *six6* genes (i.e., *SIX4-like*, *SIX1B*, and *SIX6*), as well as *RTN1A-like* (Figure 6A; full annotation information in Additional File S3). The large region contained many other genes in this region, including *fibroblast growth factor receptor-like 1* (56.48-56.56 Mbp), as well as *homeobox protein OTX2* (57.88-57.89) and three genes annotated as *60 kDa lysophospholipase* (also termed *asparaginase*; *ASPG*) within the region with the most conserved homozygosity (Figure 4B).

**Figure 6.**
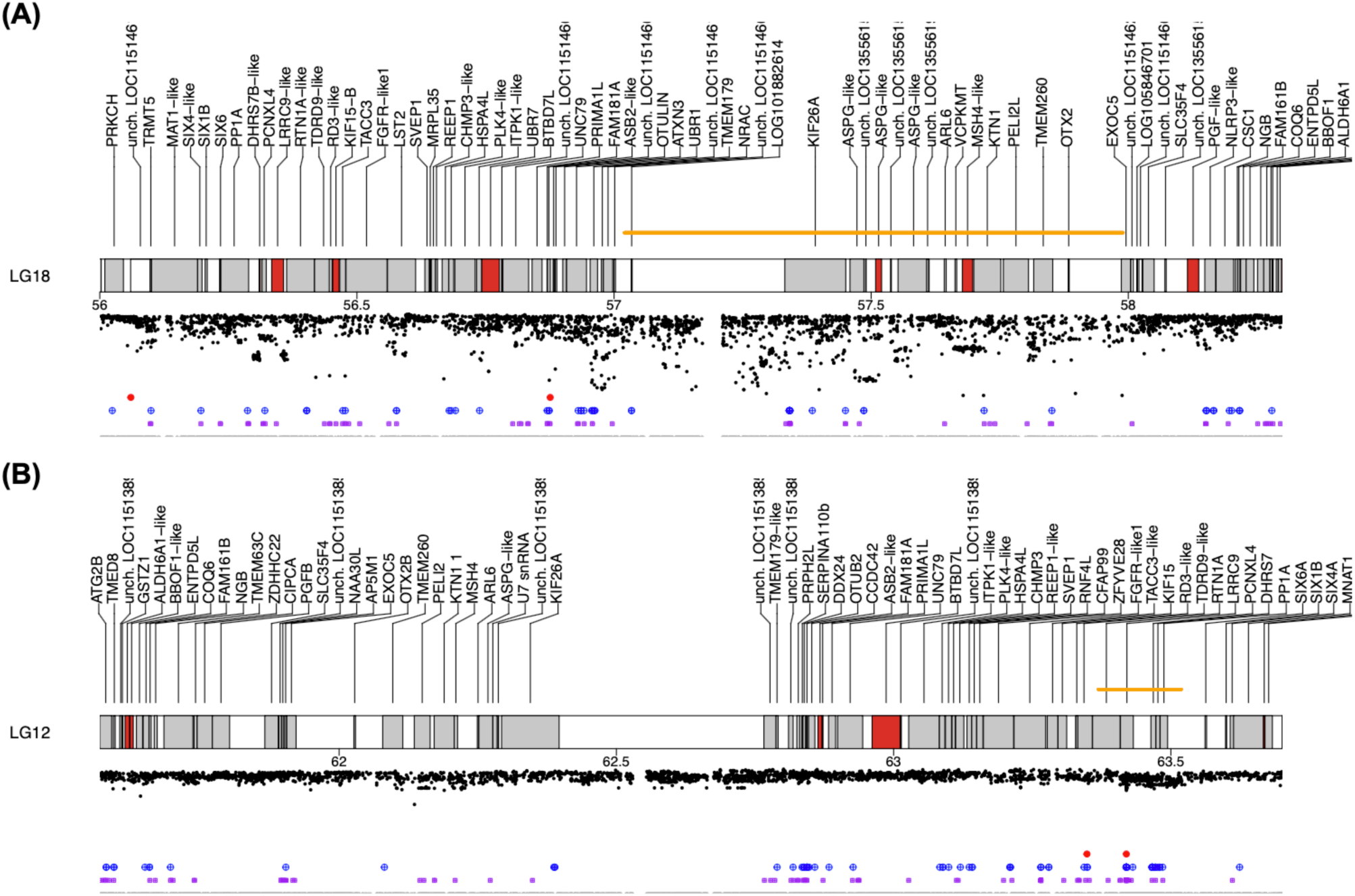
Chr18 region of interest from 56.0-58.3 Mbp with annotated genes and per-locus F_ST_ between Early Stuart and Stuart-Summer populations (shown in negative-oriented axis). Banding shows whether the gene is protein coding (grey) or is a putative pseudogene (red). Orange bars indicate regions of high linkage in Chr18 (57.02-57.99 Mbp) and Chr12 (63.37-63.52 Mbp). At the bottom of the plot are the positions for missense mutations (red dots), and high (blue) or moderate (purple) effect mutations, as well as modifier SNPs (grey dots), as defined by snpEff. Full gene names for the regions and heatmaps including flanking regions are shown in Additional Files.

The island of divergence held 3,791 SNPs, of which the impact was estimated as high (n = 2; 0.1%), moderate (n = 60; 1.6%), or low (n = 71; 1.9%) by snpEff (see Data Availability). These values were very similar to that at a chromosome-wide scale for Chr18, suggesting that the region was not enriched for SNPs of specific effect (Table S1). The estimated high effect SNPs were at positions 56,060,386 bp and 56,875,890 bp, and their expected effects included splice site disruption (splice acceptor variant & intron variant) within a long non-coding RNA (LOC115146164) and transcription start site loss in a gene annotated as *ankyrin repeat and SOCS box protein-2 like* (Table S2). The latter SNP has highest prevalence in the Chilcotin and Quesnel regions (Table S2), and does not segregate between the Stuart Watershed run timing groups (Table S2). A lack of difference in the Nechako makes this SNP therefore not relevant in understanding the differences between Stuart-Summer and Early Stuart at the island of divergence on Chr18.

### Homeologous region on Chr12 containing a known island of divergence

The homeologous region to the above Chr18 island of divergence on Chr12 was inspected (63.7 – 61.6 Mbp; chromosome assembly orientations are opposite), which included the gene region found to have broad geographic association to life history traits or run timing, *lrrc9* (e.g., Veale and Russello 2017; Tigano and Russello 2022; Barry et al. 2025). An unclustered heatmap showing genotypes for all samples ordered by reporting unit shows a region of strong linkage from 63.37 – 63.52 Mbp (with a short region with broken linkage at 63.42 Mbp; i.e., within the *lrrc9* gene) including strong linkage across the expected region from previous studies (Figure 7; Additional File S4 for full region). However, this region did not show a signature of selection for the populations inspected in the extended haplotype homozygosity analysis (Figure 5B).

**Figure 7.**
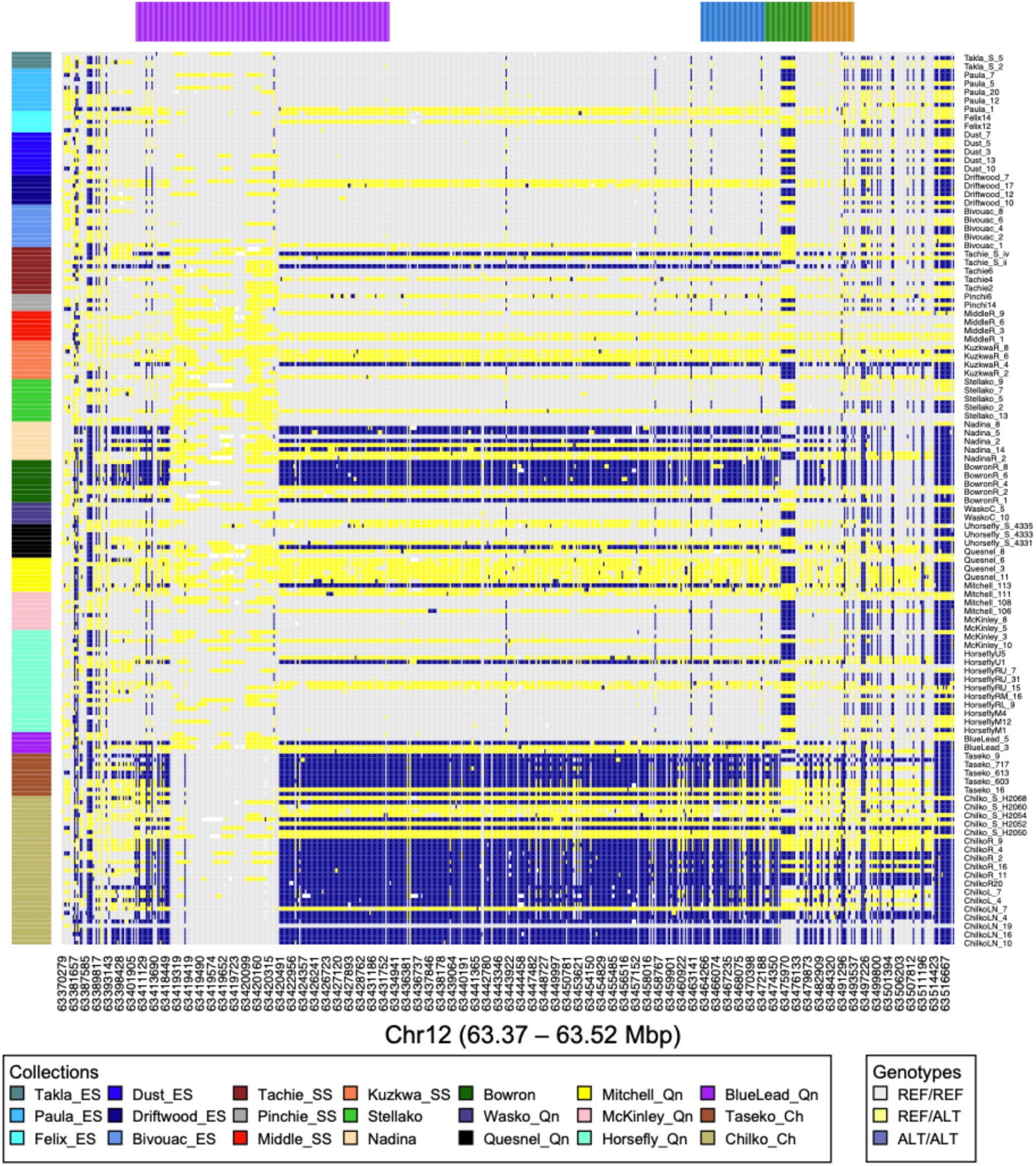
Region of Chr 12 with high linkage (63.37 – 63.52 Mbp) contained within the homeologous region to the Chr18 island of divergence. Region contains the SNP identified by Veale and Russello (2017) associated with life history traits, as well as genes *lrrc9* (top colours: purple), *pcnxl4* (blue), *dhrs7* (green), and *ppp1ca* (orange). A high impact SNP with potential splice junction disruption present at 63,419,652 bp (within *lrrc9*). Sample collection site colours are shown along left vertical axis. Acronyms (regions) in collections as per Figure 2 legend.

Most populations had multiple haplotype alleles at the haploblock (Table 2). The homozygous alternate haploblocks (representing the beach/late phenotype; see *Methods*) were most prevalent in the Early summer runs from Bowron River (70% of samples) and Chilcotin region (Taseko; 90%) as well as the Summer run in the Chilcotin region (Chilko; 74%). Interestingly, the homozygous alternate haploblock (beach/late) was rare in the Nechako region (i.e., Early Stuart = 0%; Stuart-Summer = 10%; Stellako (Summer) = 0%), except in the Nadina River (44%). There did not appear to be a clear association with population-level run timing for the Chr12 haploblock in these upper Fraser River samples.

The homeologous region on Chr12 (63.7 – 61.6 Mbp) contains genes of interest largely consistent with Chr18, including *reticulon-1A* RTN1A (63.359 – 63.409 Mbp), *lrrc9* (63.409 – 63.432 Mbp), *six6a* (63.562 – 63.564 Mbp), and *six4a* (63.607 – 63.614 Mbp; Figure 6B). The region of high LD (63.37 – 63.52 Mbp) begins near *lrrc9* (Figure 7; Additional File S4), although linkage breaks down within a short punctuated section within *lrrc9* (described above).

The snpEff analysis of the region on Chr12 had a similar frequency profile of SNP effects to Chr18 (Table S1). Two high impact SNPs were observed at positions 63,348,642 bp and 63,419,652 bp, a potential splice site disruption SNP (splice acceptor variant & intron variant) within a “low quality protein” *ATP-dependent RNA helicase TDRD9-like* and a splice donor variant & intron variant within *lrrc9*, respectively. The latter SNP is found throughout the sampling range (Table S2) and is within the splice site at the 3’ end of exon 14. There were no homozygous alternate genotypes observed for this SNP (Table S2). Inspecting NCBI RNA-seq expression tracks for *lrrc9* (NCBI Gene ID: 115138988) exon 14 through exon 15 including the intron shows 10 samples with expression of the intron (of 106 samples total), which was unusual for other introns in the gene. The tissues showing reads mapping to the intron were from female (11-month-old) upper jaw, ovaries, and pituitary, as well as unknown sex smolts in the head kidney.

### Metapopulation differentiation in Early Stuart sockeye

To determine whether there were any punctuated genomic regions of differentiation and to reassess genome-wide differentiation within the Early Stuart metapopulation, a dataset including the four populations with the greatest sample sizes was analyzed (n = 37 samples, 611,155 SNPs). This included four collections from throughout the two-lake system (Figure 1): Driftwood River, northern end of Takla Lake (n = 7); Dust Creek, western arm of Takla Lake (n = 10); Bivouac Creek, southern end of Takla Lake (n = 10); and Paula Creek, western end of Trembleur Lake (n = 10; Table 1).

Most pairwise Early Stuart comparisons had *F*_ST_ 95% confidence intervals (CI) that overlapped zero, suggesting no discernable differentiation (Table S3). Two comparisons had very low, but non-zero *F*_ST_ values: Driftwood River vs. Bivouac Creek (95% CI *F*_ST_ of 0.0017-0.0021; mean = 0.0019); and Driftwood River vs. Paula Creek (95% CI: 0.0008-0.0012; mean = 0.0010). Furthermore, a PCA shows a general overlap of all ellipses around each collection (Figure S4A), and a DAPC recovers similar trends to *F*_ST_ pairwise comparisons, where Driftwood is projected furthest from Bivouac Creek and Paula Creek, with some overlap between Bivouac Creek and Driftwood River (Figure S4B). Therefore, the geographically most distant locations (i.e., Paula Creek or Bivouac Creek vs. Driftwood River) had the greatest genetic difference, albeit even this was very low. Importantly, there were no detected genomic regions with punctuated elevated differentiation between the Early Stuart collection sites (Figure S4B).

To put the slight non-zero genetic distance observed within the metapopulation into context, contrasts between an Early Stuart collection (Paula Creek) and populations from outside the Early Stuart group were conducted. Paula Creek compared with Kuzkwa River (Stuart-Summer) shows an average genome-wide *F*_ST_ of 0.0091 (95% CI: 0.0089-0.0093) based on this dataset, which is 9.1x higher than Driftwood River vs. Paula Creek. Paula Creek compared with Bowron River shows an average genome-wide *F*_ST_ of 0.0355 (95% CI: 0.0353-0.0358), or 35.5x higher than Driftwood vs. Paula. It is also of note that the Driftwood collection had the lowest sample size of the four collections from the Early Stuart, which as described above has resulted in collections being positioned slightly differently in the genetic dendrogram and could affect the reliability of its estimated allele frequencies.

## Discussion

The main finding of this study is the identification of a large island of divergence putatively under selection in the Stuart-Summer, Nadina (Early summer), and Stellako (Summer) sockeye. This is particularly significant because the haploblock is in a homeologous region to a known island of divergence on Chr12 that has been associated with life history and run timing variation throughout the species range, including within the Fraser River Watershed (Veale and Russello 2017; Euclide et al. 2024; Barry et al. 2025). It is interesting to note that the Chr12 haploblock has very low frequency of the putative beach/late haploblock throughout most of the region where the Chr18 haploblock presence is segregating between run timing groups (i.e., Early Stuart vs. Stuart-Summer, Nadina and Stellako). It is possible that the Chr18 haploblock could have a role in run timing variation in populations without run timing-related variation within the Chr12 haploblock. However, Nadina sockeye exhibit selection signatures on the Chr18 haploblock but also have both alleles of Chr12. Given that there was no evidence for selection within the Chr18 region in other populations, it suggests that selection has acted specifically upon Stuart-Summer, Nadina, and Stellako sockeye. Whether the Chr18 region is affected by structural variation such as an inversion impeding recombination (Wellenreuther and Bernatchez 2018), and whether this structural variation may differ between populations, is unknown and merits investigation with long-read technology.

Run timing is an important phenotype contributing to the portfolio effect in salmon populations (Schindler et al. 2010). It may allow stocks to time their spawning and incubation periods appropriately for stream temperatures to provide offspring with the best chance at survival (Lisi et al. 2013). Empirical data on spawning stream temperatures suggests that Early Stuart streams are colder than the Stuart-Summer spawning grounds (e.g., Forfar Creek being significantly colder than Tachie River from March through December, *pers. comm.* Andrea Stirling). Takla monitoring has found this on most Early Stuart streams on Takla Lake (small tributaries), with the larger tributaries in Stuart-Summer spawning grounds (lake outlet rivers) being warmer and where sockeye spawning must occur later to avoid developing and emerging as fry in winter rather than spring (Cory Williamson and Takla Nation, *pers. obs.*). Although all four Nechako stocks would be considered stream spawners (not beach), the Early Stuart are somewhat unique in their spawning habitat of smaller, snowmelt drive streams rather than larger lake-influenced streams (i.e., Stuart-Summer, Nadina, and Stellako). This temperature difference fits with reasoning for the earlier arrival time and provides an interesting contrast between two separate forms of the Chr18 haploblock (Early Stuart vs. Stuart-Summer, Nadina, Stellako). Genetic variations regulating migration or spawn times would allow populations to adapt to these different stream conditions.

In many salmonids there are specific genomic regions that have underwent independent evolution pathways for this trait. As demonstrated here and in coho salmon and chum salmon by Barry et al., (2025), it appears that this pathway may in some cases involve either homeologous chromosome at the *lrrc9* or *lrrc9-like* regions, although both show different characteristics in sockeye salmon. Chr12 shows a consistent haploblock presence and long-term conservation throughout the species range with a relatively small haploblock holding *lrrc9.* Chr18 has thus far only been observed in the Nechako region (in sockeye), and largely where the Chr12 haploblock is not widely variable. It is interesting to note that the Early Stuart metapopulation does not hold a conserved haploblock on Chr18, and so the adaptive variation separating the two proximal run timing groups (Early Stuart vs. Stuart-Summer) may be related to delaying the return for the Stuart-Summer group, if the region is indeed involved in modulating run timing. Given that the *lrrc9* region is being proposed as a putative ‘master regulatory region’ for run timing in other salmonids (Barry et al. 2025), it is reasonable to consider a potential role in the Chr18 region affecting run timing differences between the Stuart-Summer and the Early Stuart populations. Other explanations could exist for the Chr18 haploblock. It may still have a functional role but not in run timing, or it could be not functional and could rather be related to other reasons of island of divergence formation such as reduced recombination due to structural variation and allele fixation through drift (Renaut et al. 2013; Quilodrán et al. 2020). However, although smaller islands may form by drift during species divergence, large genomic islands rather require a combination of divergent selection and strong linkage (Quilodrán et al. 2020), and the Chr18 island would likely be classified as a large genomic island.

The Chr12 haploblock association to run timing in the present study is unclear; the haploblock is clearly present and conserved as a limited number of haplotypes, but the known ‘stream/early’ (REF/REF or REF/ALT) and ‘beach/late’ (ALT/ALT) genotypes do not correspond well to the run timings of populations assessed here (e.g., the highest frequency of the ‘late’ allele is in many early summer runs such as Taseko River and Bowron River). The Chr12 *lrrc9* haploblock was also associated with other life history variation (shore or stream spawning) throughout the range in sockeye (Veale and Russello 2017; Tigano and Russello 2022), but this is often also confounded with run timing (Barry et al. 2024). It is also worthwhile noting that the Early Stuart metapopulation has a relatively large population size (and presumably large effective population size) relative to other populations in the region, potentially affecting either the effect of drift or of selection in the emergence of islands of divergence (Renaut et al. 2013; Quilodrán et al. 2020). The detection of a SNP disrupting a splice site within *lrrc9* is notable and worthwhile following up with future transcriptome analyses focused on alternate splicing in relation to the genotype at this locus. This is particularly interesting given the complete lack of homozygous alternate genotypes at this locus, and therefore potential of a detrimental effect of the variant when homozygous.

Tracking consistent chromosome identities within and between species is facilitated by using the ancestral naming convention outlined by Sutherland et al. (2016) to support analyses related to the post-WGD state of salmonid genomes using northern pike chromosomes, which are expected to be similar to preduplicated salmonid form (Rondeau et al. 2014). The haploblock on Chr18 (56.3 – 58.0 Mbp; present study) is within the chromosome arm corresponding to the ancestral chromosome 15.2 (see Sutherland et al. 2016). The haploblock on Chr12 of sockeye and Chr10 of pink salmon associated with run timing in Alaska (Barry et al. 2025) corresponds in both species to the ancestral chromosome 15.1. Here we observed the haploblock on Chr12 with strong linkage at 63.37 – 63.52 Mbp. The observation of both homeologs showing islands of divergence within a single species was also noted by Barry et al. (2025), who found secondary regions of high differentiation in ancestral 15.2 in chum and coho salmon. However, 15.2 has not yet been identified as having an island of divergence at the *lrrc9-like* region in sockeye salmon, and is the first time this region has been shown to exhibit evidence of selection.

Based on evolutionary theory, resolution of WGD state typically involves subfunctionalization, neofunctionalization, or elimination of one ohnolog to maintain effective dosage for gene network function (Crow and Wagner 2006). The observation that *lrrc9-like* is a putative pseudogene may provide additional information about the potential mechanisms underlying the variation. Barry et al. (2025) propose that the region of differentiation is likely driving phenotypic effects through transcription regulation activity rather than changes in protein coding sequences due to the lack of non-synonymous mutations identified and the prevalence of transcription factors in the region. The splice site variant observed in the present work on Chr12 is of note in terms of potential drivers of phenotypic differences, and merits further analysis. If *lrrc9-like* is a pseudogene (as it appears to be), the finding of an island of divergence under selection on Chr18 would support the lack of direct involvement of this gene in any downstream phenotype. There are other genes in the major differentiated region identified here in sockeye salmon that may be of interest regarding a migratory timing phenotype, such as the *six6* genes and *homeobox protein OTX2* (*orthodenticle homeobox 2*). Along with *vgll3*, proximal *six6* genes have been linked to age-at-maturity in Atlantic salmon (Barson et al. 2015; Sinclair-Waters et al. 2022). *OTX2* is a transcription factor involved in brain and sensory organ development (Beby and Lamonerie 2013). *OTX2* is involved in the formation of the pituitary gland in zebrafish (Bando et al. 2020), and the pituitary-gonadal axis activity in salmonids is connected to season and directly related to homing adults, return migration and spawning preparation (Onuma et al. 2009; Ueda 2011). Furthermore, the role of *OTX2* in the development of olfactory system, including imprinting and homing has been demonstrated in the anemonefish *Amphiprion percula* (Veilleux et al. 2013), of interest given the important role of the olfactory system in homing migration in sockeye salmon (Bett et al. 2018).

The present study therefore identifies new evidence that points to potential involvement of the proposed master regulatory region on ancestral chromosome 15 (Chr18) that has taken an alternate evolutionary path in the alternate homeologous chromosome path to that previously identified in sockeye salmon (Tigano and Russello 2022; Barry et al. 2025). This supports the conclusions of Barry et al. (2025), as well as underscoring the unique genetics and evolutionarily significant adaptation route of upper Fraser River sockeye salmon. As experimental validation of any functions of the island of divergence on Chr18 (or Chr12) have not been conducted, the role of this region in run timing remains putative. Although other phenotypic run timing differences occur in the populations analyzed in this study, for example the two run timings of the Chilcotin Watershed sockeye (i.e., Chilko as Summer and Taseko as Early Summer), there were no similar regions of major differentiation observed in these other populations using the available samples. It should also be noted that within population variation in run timing was not possible with the samples present in the current dataset, as this was not part of sample collection goals or metadata. Potential intrapopulation effects of the haploblock on Chr12 are therefore not possible to explore here.

Considering the potentially important roles of ancestral chromosome 15.1 and 15.2, it is worth briefly noting other features of these chromosomes in the evolution of salmonids. Both chromosome segments are part of ancestral and conserved fusion events; ancestral 15.1 is proposed to have been fused to ancestral 4.1 prior to the divergence of genera *Salmo* and *Oncorhynchus*, although lineage-specific fissions and fusions occurred in the lineage leading to the pink, chum, sockeye branch (Sutherland et al. 2016). Ancestral 15.2 was part of a conserved fusion to ancestral 25.2 that occurred prior to the diversification of *Oncorhynchus*; this fusion remains in all *Oncorhynchus* species. Another interesting feature related to ancestral 15 is that, chromosome arm 15.1 is within the chromosome that holds the master sex determining gene in Arctic charr *Salvelinus alpinus* and brook charr *S. fontinalis* (Sutherland et al. 2017). A continued understanding of salmonid chromosome evolution is likely to yield valuable insights in the evolution of phenotypic variation in the salmonids. This work also points to the high value in ensuring homeologous and orthologous definitions for all salmonid chromosomes are considered in comparative genomic studies involving major effect loci.

A final note on the Early Stuart metapopulation is worthwhile to state here: we did not identify any punctuated regions of differentiation between streams, and therefore the previous findings of low to negligible differentiation with higher sample sizes but lower marker density (i.e., Beacham et al. 2004; Rondeau 2022), should be sufficiently reliable in this conclusion. The slight increase but still near-zero differentiation at greater distances is interesting but would benefit from larger sample sizes on all populations. It is possible that any phenotypic variation observed between streams could be associated with plasticity, environmental effects, or epigenetic differences between populations (Lamka et al. 2022). Therefore, these results support previous work indicating an inability to discriminate between populations using genetic stock identification (GSI) and support the connectivity and characteristics of a metapopulation (Bradford and Braun 2021), but do provide some evidence for slight spatial differentiation that is worthwhile considering as not completely panmictic. Although low sample size appears to have resulted in misplaced populations in the genetic dendrogram (most noticeable with sample size under n = 5), and therefore raises the potential concern that this could drive the non-zero *F*_ST_ observed between Driftwood and other more distant streams, the observation that Driftwood (n = 7) compared to Dust Creek (n = 10), the more proximal sampling location, had an estimated zero differentiation indicates that the conclusion of no differentiation could be estimated at this sample size. It should also be noted that the samples used to represent the Early Stuart sockeye in this study were from 2005, and if large scale genetic changes occurred for salmon spawning in these tributaries since 2005, these results may not reflect current day genetic signatures. Further investigation of the extent of gene flow in this metapopulation could be conducted by parentage-based tagging (PBT) or other tagging methods that track returning adults to determine the extent of flow between tributaries. Epigenetic analyses could be conducted to determine the level of differentiation epigenetically between streams with different environmental parameters (e.g., Venney et al. 2021; Lamka et al. 2022).

## Conclusions

In the present study we made use of existing genomic resources to characterize upper Fraser River sockeye salmon populations, and the findings of this study would not have been possible without the generation and publication of large-scale, open-source genomic resources and metadata. The island of divergence observed on Chr18 (ancestral 15.2) appears to be under selection only within the Stuart-Summer/ Nadina/ Stellako group, and given the role observed for the homeologous region on Chr12, may be involved in adaptive variation underlying run timing, and in differentiating run timing from the nearby Early Stuart sockeye metapopulation. The Chr18 (15.2) region identified here provides further evidence for the importance of this genomic region in both homeologous chromosomes in different populations of the same species, evolving independently in multiple species. This region and its gene content, patterns of gene expression including alternate splicing, and the potential involvement of structural variation heterogeneity at Chr18 in the upper Fraser River and elsewhere merits further investigation.

## Supporting information

Supplemental Results

## Acknowledgements

Thanks to Drs. Ben Koop, Eric Rondeau, Wes Larson, and Kris Christensen for discussion on the major differentiation regions, and to Linda Stevens of UFFCA for project support and comments on the manuscript. Thanks to the Upper Fraser Fisheries Conservation Alliance (UFFCA) for supporting this work. Thanks to four anonymous reviewers and the Editor for comments on an earlier manuscript draft.

## Data Availability

SNP data were downloaded from FigShare: https://doi.org/10.25387/g3.25705428.v1 The following analytic repositories supported this analysis:

Code and README for project: https://github.com/bensutherland/ms_oner_estu Population genetic functions: https://github.com/bensutherland/simple_pop_stats Predicted effects of polymorphisms by snpEff in Chr18 and Chr12 available on FigShare: https://doi.org/10.6084/m9.figshare.32885771.

## Additional Materials

A Supplemental Results section is provided with supplemental figures and tables.

**Additional File S1.** Genotypes of top differentiated loci in major differentiation region.

**Additional File S2.** Heatmap of Chr18 region of interest in all samples, sorted by reporting unit.

**Additional File S3.** Annotated gene content in major differentiation region.

**Additional File S4.** Heatmap of Chr12 region of interest in all samples, sorted by reporting unit.

