## Supplemental Results for "Genomic islands of divergence reveal selection on the alternate homeolog in upper Fraser River sockeye run-timing groups"

**Table of Contents:**


| | |
|---|---|
| Figure S1. Genetic dendrogram, all populations | Page 2 |
| Figure S2. Chilcotin PCA, DAPC, and loadings | Page 3 |
| Figure S3. Quesnel PCA, DAPC, and loadings | Page 4 |
| Figure S4. Early Stuart PCA, DAPC, and loadings | Page 5 |
| Figure S5. RNA-seq read coverage *lrrc9* and *lrrc9-like* | Page 6 |
| Table S1. Summary of snpEff predicted impacts per chromosome and region | Page 7 |
| Table S2. Genotype frequencies of snpEff loci of interest | Page 7 |
| Table S3. $F_{ST}$ table with focus on Early Stuart collections | Page 7 |

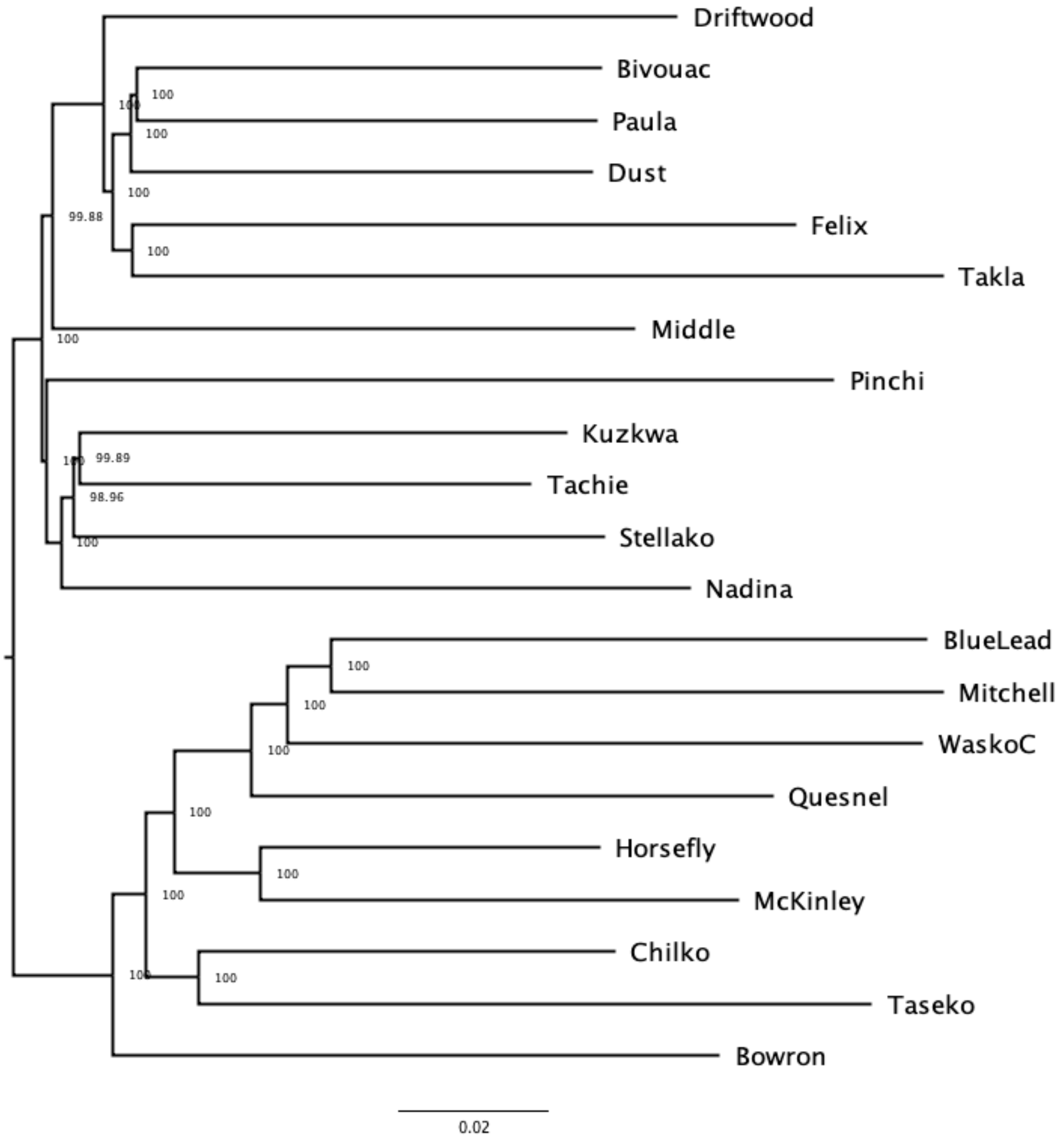


**Figure S1.** Genetic dendrogram with midpoint root constructed using Weir-Cockerham $F_{ST}$ values. Includes all collections from the whole-genome resequencing dataset using the LD-filtered SNPs.

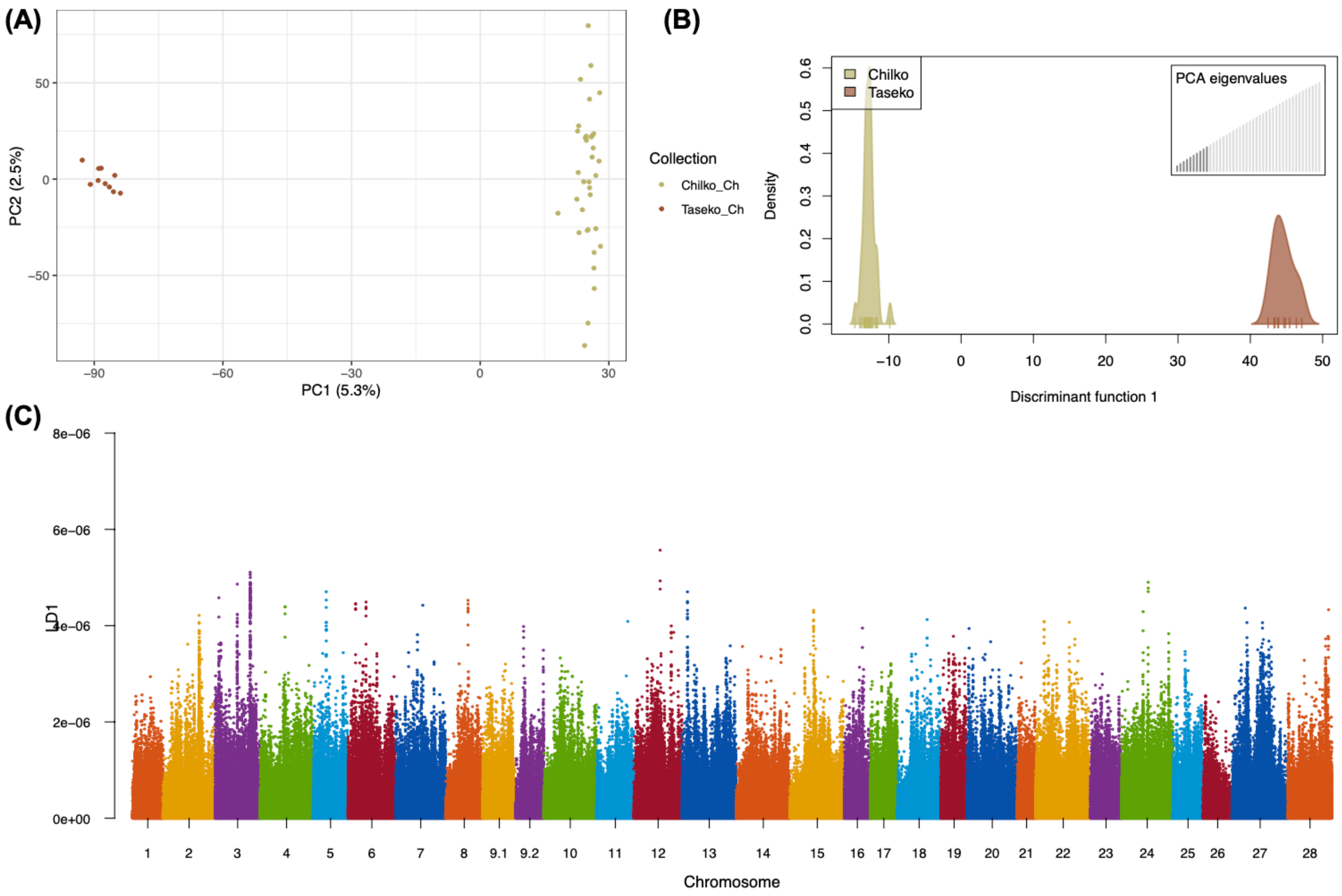


**Figure S2.** Chilcotin River Watershed **(A)** PCA; **(B)** DAPC; and **(C)** DAPC loadings on chromosomes.

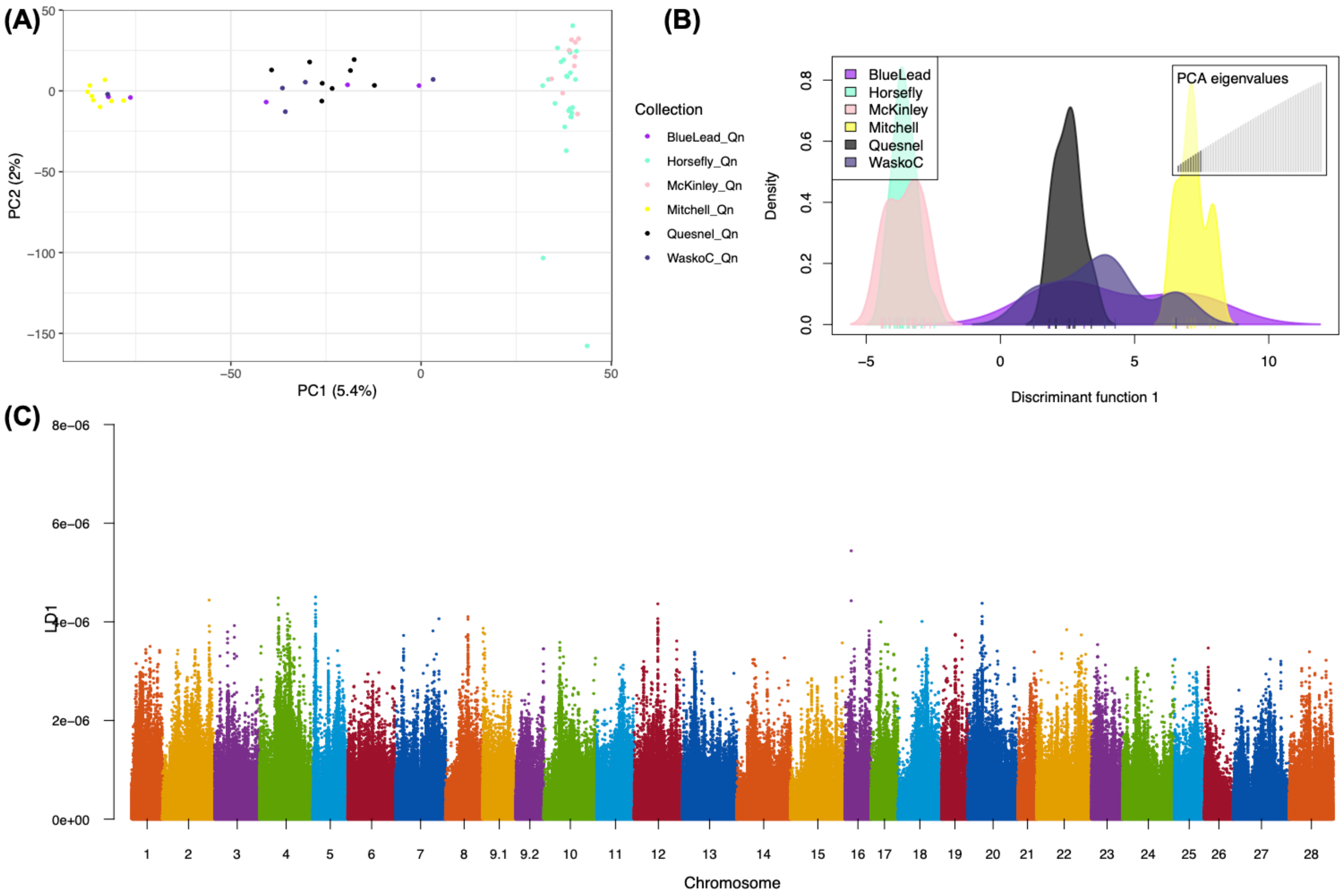


**Figure S3.** Quesnel and Horsefly River system **(A)** PCA; **(B)** DAPC; and **(C)** DAPC loadings on chromosomes.

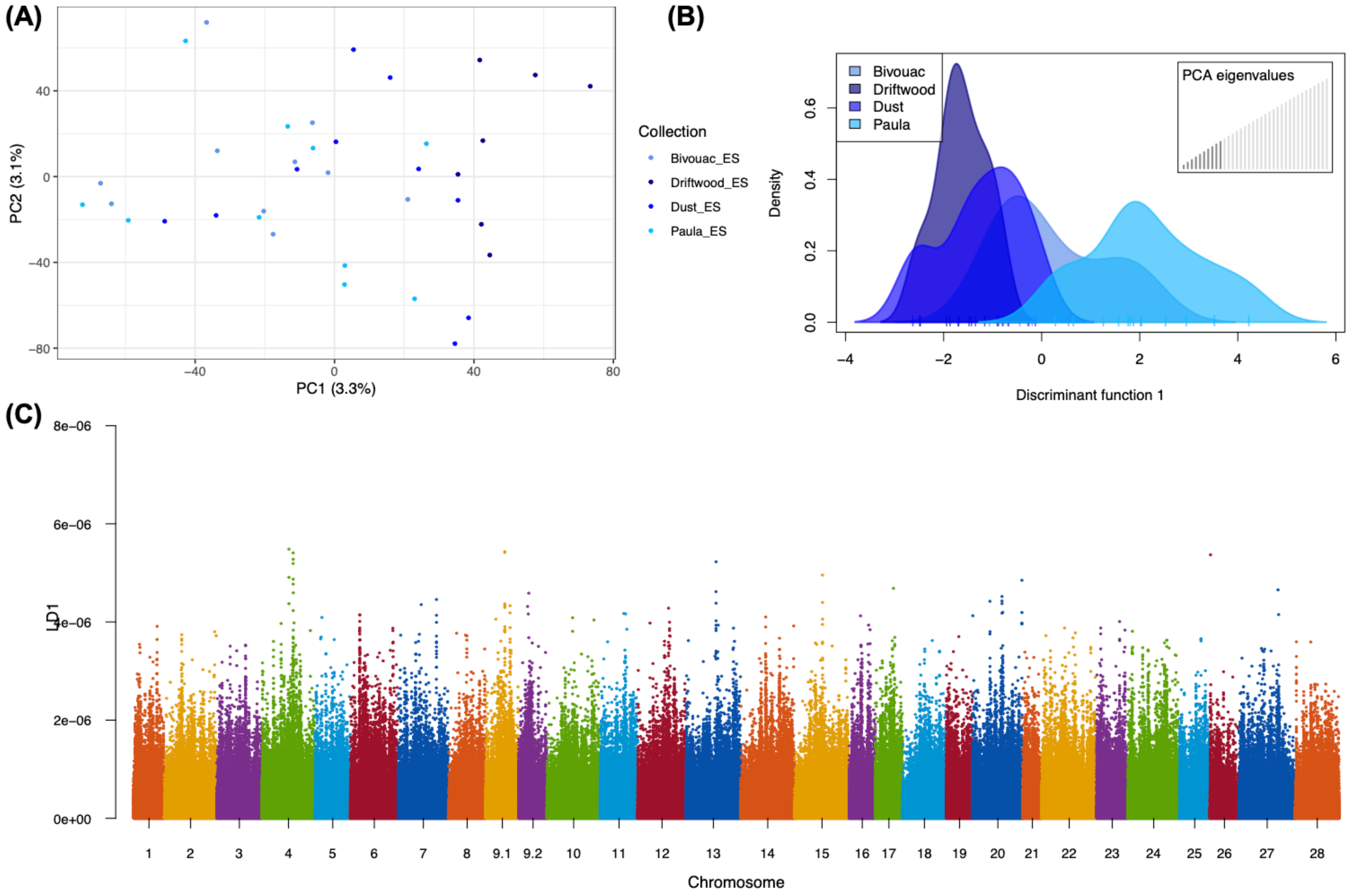


**Figure S4.** Early Stuart **(A)** PCA; **(B)** DAPC; and (C) DAPC loadings on chromosomes. Only includes collections with at least five samples.

**(A)** *lrrc9* on chr12

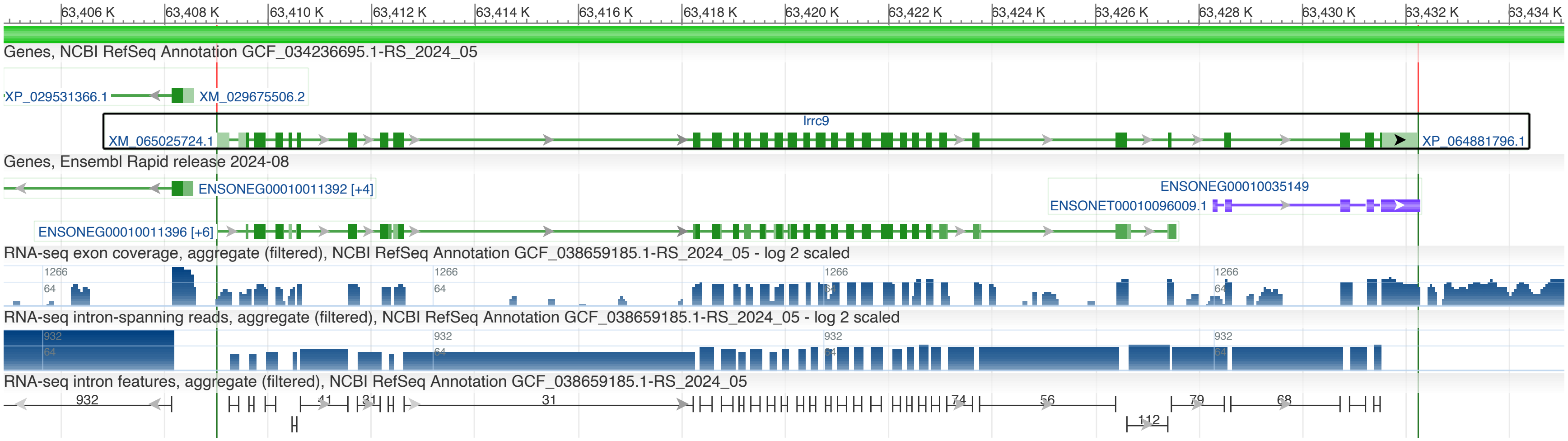


**(B)** *lrrc9-like* on chr18

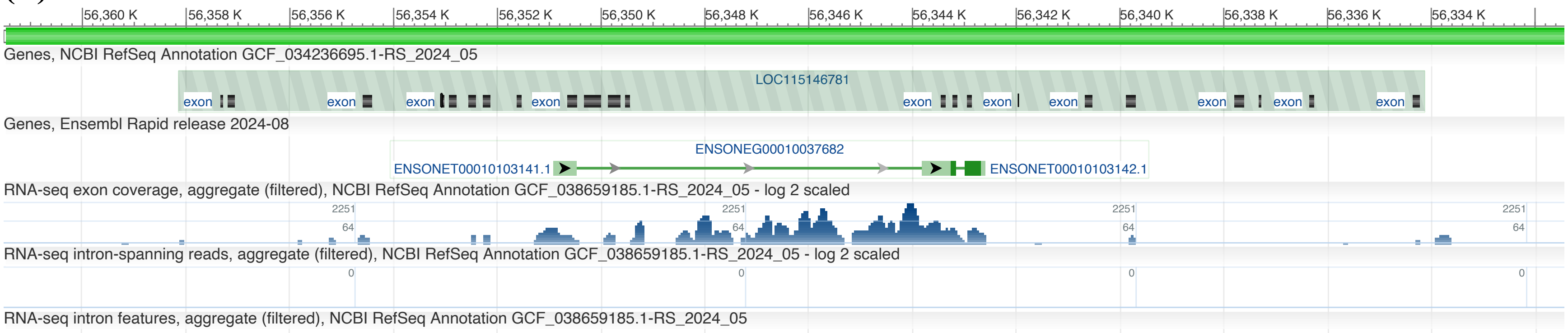


**Figure S5.** Predicted exons and RNA-seq coverage (aggregate) from NCBI Genome Viewer for (A) *lrrc9*; and (B) *lrrc9-like* (predicted pseudogene).

**Table S1.** SNP effects on Chr18 and Chr12 as predicted by snpEff.

| **Chr** | **Region** | **SNPs (n)** | **High** | | **Moderate** | | **Low** | | **Modifier** | |
|---|---|---|---|---|---|---|---|---|---|---|
| 18 | All | 144,025 | 131 | 0.1% | 2,185 | 1.5% | 2,493 | 1.7% | 139,216 | 96.7% |
| 18 | 56.0-58.3 Mbp | 3,791 | 2 | 0.1% | 60 | 1.6% | 71 | 1.9% | 3,658 | 96.5% |
| 12 | All | 139,774 | 99 | 0.1% | 1,882 | 1.3% | 1,946 | 1.4% | 135,847 | 97.2% |
| 12 | 61.6-63.7 Mbp | 3,885 | 2 | 0.1% | 84 | 2.2% | 98 | 2.5% | 3,701 | 95.3% |

**Table S2.** Loci of interest from snpEff analysis for chr12 and chr18 showing genotype frequencies per collection.

| | | | | Splice site variant, *lrrc9* NC_088407.1 (chr12), 63,419,652 bp | | | Start site lost, *asb2* NC_088413.1 (chr18), 56,875,890 bp | | |
|---|---|---|---|---|---|---|---|---|---|
| **Region** | **Repunit** | **Collection** | **n** | **REF/ REF** | **REF/ ALT** | **ALT/ ALT** | **REF/ REF** | **REF/ ALT** | **ALT/ ALT** |
| Stuart | Early-Stuart | Bivouac | 10 | 9 | 1 | | 10 | | |
| | | Driftwood | 7 | 6 | 1 | | 7 | | |
| | | Dust | 10 | 8 | 2 | | 10 | | |
| | | Felix | 5 | 5 | | | 5 | | |
| | | Paula | 10 | 8 | 2 | | 9 | 1 | |
| | | Takla | 4 | 3 | 1 | | 4 | | |
| | Summer | Kuzkwa | 9 | 5 | 4 | | 9 | | |
| | | Middle | 7 | 1 | 6 | | 7 | | |
| | | Pinchi | 4 | 3 | 1 | | 4 | | |
| | | Tachie | 11 | 6 | 5 | | 11 | | |
| Nadina/Francois | Early summer | Nadina | 9 | 3 | 4 | | 9 | | |
| Francois/ Fraser | Summer | Stellako | 10 | 3 | 6 | | 9 | 1 | |
| Bowron | Early summer | Bowron | 10 | 8 | 2 | | 10 | | |
| Quesnel | Summer | BlueLead | 5 | 2 | 3 | | | 4 | 1 |
| | | McKinley | 9 | 5 | 4 | | 7 | 2 | |
| | | Mitchell | 8 | 2 | 6 | | | 4 | 4 |
| | | Wasko | 5 | 3 | 2 | | 1 | 3 | 1 |
| | | Horsefly* | 24 | 18 | 6 | | 17 | 7 | |
| | | Quesnel | 8 | 1 | 6 | | 2 | 3 | 3 |
| Chilcotin | Early Summer | Taseko | 10 | 10 | | | 8 | 2 | |
| | Summer | Chilko* | 35 | 30 | 4 | | 21 | 10 | 4 |

*Missing data present for Nadina (x2), Stellako (x1), Quesnel (x1), and Chilko (x1) for lrrc9 variant.*

**Table S3. (A)** Average $F_{ST}$ between Early Stuart collections showing non-weighted estimate in the bottom and weighted estimate in the top section; or **(B)** 95% confidence interval of $F_{ST}$ showing the upper limit in the top and lower limit in the bottom.

**(A)**

| | **Bivouac** | **Driftwood** | **Dust** | **Paula** |
|---|---|---|---|---|
| **Bivouac** | NA | 0.0056 | 0.0010 | 0.0008 |
| **Driftwood** | 0.0019 | NA | 0.0022 | 0.0046 |
| **Dust** | -0.0011 | -0.0009 | NA | 0.0016 |
| **Paula** | -0.0011 | 0.0010 | -0.0006 | NA |

**(B)**

| | **Bivouac** | **Driftwood** | **Dust** | **Paula** |
|---|---|---|---|---|
| **Bivouac** | NA | 0.0021 | -0.0009 | -0.0010 |
| **Driftwood** | 0.0017 | NA | -0.0007 | 0.0012 |
| **Dust** | -0.0013 | -0.0011 | NA | -0.0004 |
| **Paula** | -0.0013 | 0.0008 | -0.0007 | NA |
