## Supplementary figures and images for "Genomic islands of divergence reveal selection on the alternate homeolog in upper Fraser River sockeye run-timing groups"

### Additional File S2

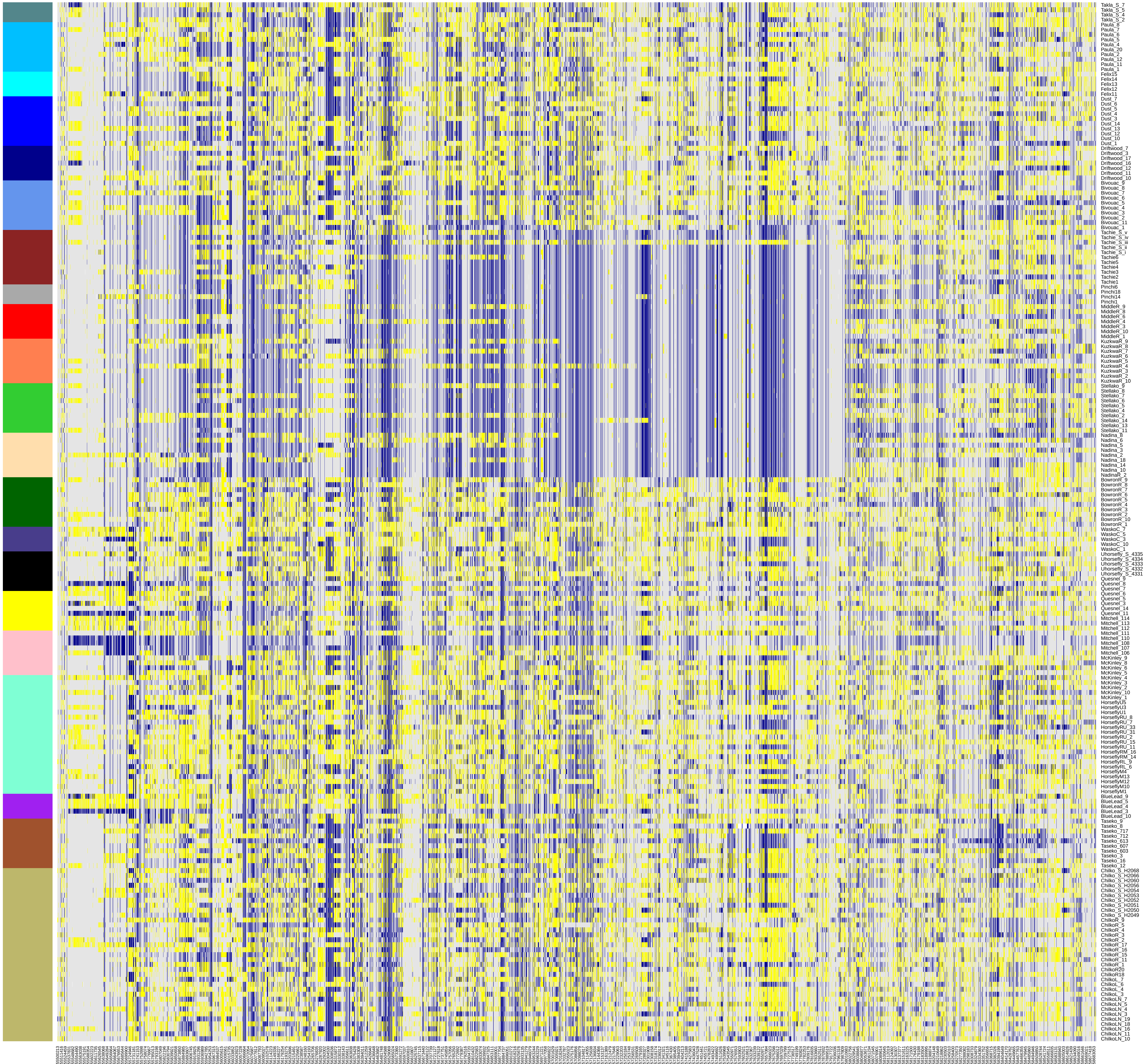

### Additional File S4

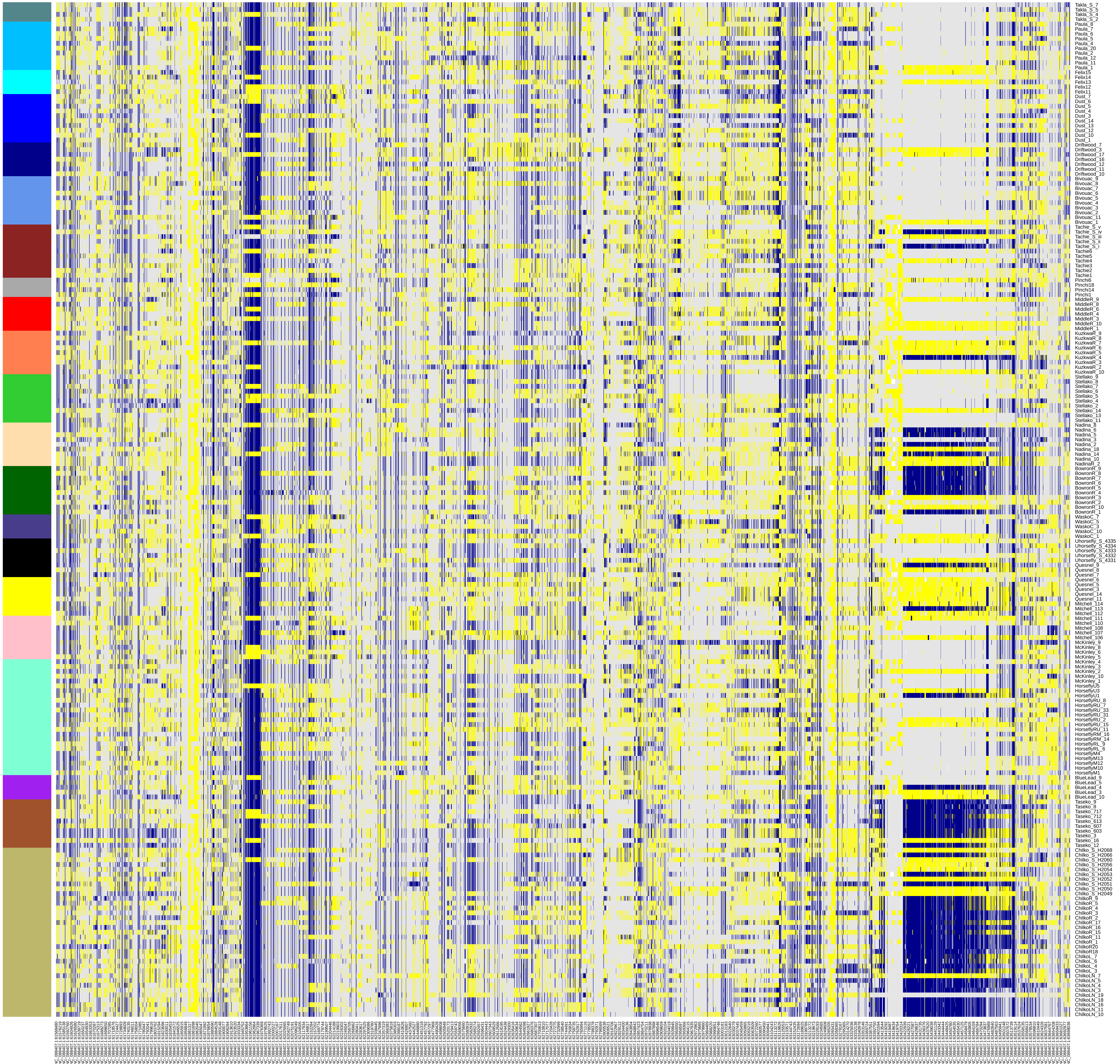
